# Reading reshapes stimulus selectivity in the visual word form area

**DOI:** 10.1101/2023.10.04.560764

**Authors:** Vassiki S. Chauhan, Krystal C. McCook, Alex L. White

## Abstract

Reading depends on a brain region known as the “visual word form area” (VWFA) in left ventral occipito-temporal cortex. This region’s function is debated because its stimulus selectivity is not absolute, it is modulated by a variety of task demands, and it is inconsistently localized. We used fMRI to characterize the combination of sensory and cognitive factors that activate word-responsive regions that we precisely localized in 16 adult humans (4 male). We then presented three types of character strings: English words, pseudowords, and unfamiliar characters with matched visual features. Participants performed three different tasks while viewing those stimuli: detecting real words, detecting color in the characters, and detecting color in the fixation mark. There were three primary findings about the VWFA’s response: (1) It preferred letter strings over unfamiliar characters even when the stimuli were ignored during the fixation task; (2) Compared to those baseline responses, engaging in the word reading task *enhanced* the response to words but *suppressed* the response to unfamiliar characters. (3) Attending to the stimuli to judge their font color had little effect on the response magnitudes. Thus, the VWFA is uniquely modulated by a cognitive signal that is specific to voluntary linguistic processing and is not additive. Functional connectivity analyses revealed that communication between the VWFA and a left frontal language area increased when the participant engaged in the linguistic task. We conclude that the VWFA is inherently selective for familiar orthography, but it falls under control of the language network when the task demands it.

**Significance Statement:** The function of the “visual word form area” (VWFA) is controversial. Some researchers emphasize its bottom-up visual selectivity for words, hence the region’s common name. Others argue that its activity is explained by feedback from regions that control attention or language. To seek clarity, we investigated what drives the VWFA: seeing words, attending visually to words, or trying to read words. None of those factors was sufficient on its own. Our results support a hybrid model: the VWFA has inherent selectivity for words, but its function is reshaped by voluntary language processing. Thus, with an integrated analysis of sensory inputs, task demands, and network connectivity, we provide some resolution to debates about this important region.

## Introduction

Over the course of history, developments in human culture have changed our environment and required our brains to adapt. A unique example is a region in the left ventral occipito-temporal cortex that, in literate individuals, is specialized for processing written words (Nobre et al., 1994; McCandliss et al., 2003; Dehaene and Cohen, 2011). This region is often termed the visual word form area (VWFA). While the VWFA’s plasticity during literacy acquisition is well established (Dehaene et al., 2015; Kubota et al., 2019), less is known about how its stimulus selectivity changes in the short term to meet task demands.

The nature of the VWFA’s functional specialization is debated. Some researchers characterize it primarily as a visual region tuned for a particular category of stimulus (McCandliss et al., 2003; Dehaene et al., 2005). It may serve as the “orthographic lexicon” that identifies familiar letter strings (Glezer et al., 2009; Dehaene and Cohen, 2011; Yeatman and White, 2021). Other researchers emphasize its integration of linguistic information across sensory modalities (Price and Devlin, 2011; Qin et al., 2021; Dȩbska et al., 2023). We contribute to this lively discussion by examining how task demands shape the VWFA’s stimulus selectivity as well as its communication with the canonical language network.

Like other areas around it, the VWFA is sensitive to visual stimulus properties. While it responds above baseline to many types of images, it prefers text (Ben-Shachar et al., 2007; Muayqil et al., 2015; White et al., 2023). Its response is affected by visual properties of words, such as their length (Woolnough et al., 2021), position (Rauschecker et al., 2012), and contrast (Kay and Yeatman, 2017). Beyond purely visual properties, the VWFA is also sensitive to statistics of letter combinations (Vinckier et al., 2007; Woolnough et al., 2021; Gagl et al., 2022), and word frequency (Kronbichler et al., 2004).

Moreover, the VWFA’s activity is influenced by attentional allocation and task demands (Mano et al., 2013). Words evoke larger responses when they are attended than ignored (Kay and Yeatman, 2017; White et al., 2019, 2023). There is also evidence for top-down language influences without visual input, such as during spoken language comprehension (Planton et al., 2019) or reading Braille (Reich et al., 2011; Striem-Amit et al., 2012). Indeed, the VWFA’s function is supported by its connections with language and attention networks (Yeatman et al., 2012; Stevens et al., 2017; Lerma-Usabiaga et al., 2018; Chen et al., 2019; Li et al., 2020).

To better understand the VWFA’s functional specialization, we used functional magnetic resonance imaging (fMRI) to examine the *interaction* of stimulus features and task demands. On each trial, we briefly flashed a character string that was either a real English word, a pronounceable pseudoword, or a string of unfamiliar “false font” characters (see **Figure 1C**). The false fonts were matched to the Latin fonts in visual features (size, number of strokes, perimetric complexity, etc.) (Vidal et al., 2017; Vildavski et al., 2022).

**Figure 1:**
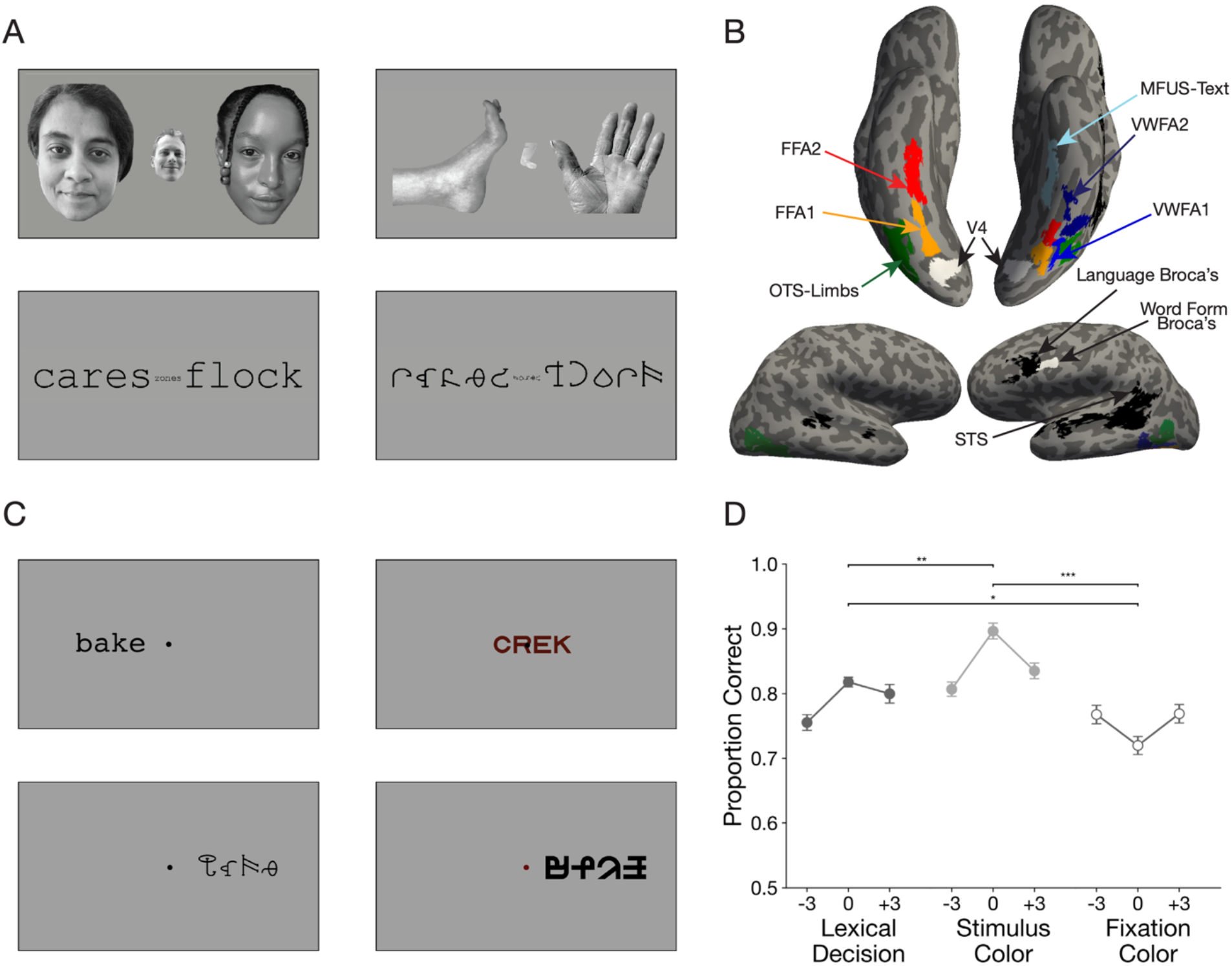
Experimental Paradigm. **(A)** Example stimuli in the visual localizer scan for category-selective regions in the ventral temporal cortex. See Extended Data Figure 1-1 for list of all stimuli used in the experiment. **(B)** Regions of interest (ROIs) in one example participant. Regions in color were defined using the localizer in panel A. Regions in black were defined from the language localizer scan, and regions in white were obtained from a previous study (White et al., 2023). See Extended Data Figure 1-2 for ROIs all individual participants. **(C)** Example stimuli in the main experiment. **(D)** Mean accuracy in the three behavioral tasks plotted as a function of the character string’s position in degrees of visual angle. Error bars are ±1 SEM. Stars and horizontal lines depict statistically significant task effects (* = p < 0.05, ** = p < 0.01, *** = p < 0.0001)

The stimulus types were randomly intermingled during three different tasks. The *fixation color task* was to report whether the fixation dot turned red. This task evaluated the VWFA’s inherent preference for words in the absence of voluntary attention or linguistic processing. The dot’s color change was simultaneous with the character string’s brief presentation and set to individual detection thresholds. The *stimulus color task* was to report whether the character string was red or gray. This task evaluated the effect of attending to the stimuli, which may result in automatic linguistic processing. The *lexical decision task* was to report whether the stimulus was a real word or not. This task required explicit linguistic processing.

By comparing stimulus responses and functional connectivity across the tasks, we ask: what maximally drives activity in the VWFA? Simply viewing words, attending to the visual properties of words, or explicitly engaging in a linguistic task?

## Materials and Methods

### Participants

We recruited 17 participants (Age range 19 to 38, 21.12 ± 4.44, 4 self-identified as male, 1 left-handed) from the Barnard College and Columbia University student body. The study was approved by the Internal Review Board at Barnard College, Columbia University. All participants provided written informed consent, acquired digitally, and were monetarily compensated for their participation. All participants had learned English before the age of five.

We chose the sample size on the basis of a power analysis on a task effect observed in a recent study with a similar experimental paradigm (White et al., 2023). Specifically, we focused on the reduced response to non-letter shapes when they were attended than when ignored (the smallest effect in that study). A simulation determined that a sample size of 17 would be sufficient to detect, with 80% power, such an effect with a predicted mean (0.037) and standard deviation (0.052, adjusted for the number of trials per condition).

To ensure high data quality, we used the following criteria for excluding functional runs and participants. If the participant moved by a distance greater than 2 voxels (4 mm) within a single run, that run was excluded from analysis. Additionally, if the participant responded in less than 50% of the trials in the main experiment, that run was removed. Finally, if half or more of the runs met any of these criteria for a single participant, that participant was dropped from the dataset. Using these constraints, the analysis reported here is based on data from 16 participants. They ranged in age from 19 to 38 years (mean = 21.12 ± 4.58,). 4 participants self-identified as male, and 1 was left-handed. A total of 6 runs were removed from three of the remaining participants due to excessive head motion.

### Equipment

We collected MRI data at the Zuckerman Institute, Columbia University, a 3T Siemens Prisma scanner and a 64-channel head coil. In each MR session, we acquired a T1 weighted structural scan, with voxels measuring 1 mm isometrically. We acquired functional data with a T2* echo planar imaging sequences with multiband echo sequencing (SMS3) for whole brain coverage. The TR was 1.5s, TE was 30 ms and the flip angle was 62°. The voxel size was 2 mm isotropic.

Stimuli were presented on an LCD screen that the participants viewed through a mirror with a viewing distance of 142 cm. The display had a resolution of 1920 by 1080 pixels, and a refresh rate of 60 Hz. We presented the stimuli using custom code written in MATLAB and the Psychophysics Toolbox (Brainard, 1997; Pelli, 1997). Throughout the scan, we recorded monocular gaze position using an SR Research Eyelink 1000 tracker. Participants responded with their right hand via three buttons on an MR-safe response pad.

### Main Experiment

#### Stimuli

**Figure 1A** shows example stimuli. A small dark fixation dot at screen center was present throughout all trials against a white background (95% of screen maximum). The dot’s diameter was 0.11 degrees of visual angle, or dva. The stimuli were character strings from three different categories: real English words (different parts of speech), pseudowords (meaningless but pronounceable strings) and visually matched false fonts. The pseudowords were generated using MCWord to match the bigram and trigram statistics of the real English words (Medler and Binder, 2005). All strings were composed of four characters each. For the full list of stimuli, see **Extended Data Figure 1-1**.

Two different fonts were used for the real words and pseudowords: Courier New and Sloan. The false fonts, BACS2 (Vidal et al., 2017) and PseudoSloan (Vildavski et al., 2022), were chosen to match the visual properties of the real fonts. The false Courier characters were matched in size, symmetry, and the number of strokes, junctions and terminations. The false Sloan characters were matched in ink area and perimetric complexity. All four fonts were scaled such that the height of the letter ‘x’ (or its false font sibling) was 0.41 dva. The distance between centers of neighboring letters was roughly 0.53 dva (ranging 0.52–0.55 across fonts). Each word was 2 dva wide, and between 0.41 and 0.78 dva tall.

On each trial, a single character string flashed for 150 ms at one of three locations: centered at fixation, 3 dva left, or 3 dva right). The stimulus was followed by a blank with only the fixation mark present for 3850 ms, during which the participant had the opportunity to respond with a button press. After every five trials, there was a rest period (no task except to fixation on the dot). The duration of the rest period was either 4, 6 or 8 s in duration (randomly and uniformly selected).

The color of the character string was dark gray on 67% of trials (70% contrast against the background) or dark red on 33% of trials. The dark red was roughly the same luminance as the dark gray. Similarly, on an independently selected 33% of trials, the fixation dot turned from dark gray to dark red during the 150 ms that the character string appeared. On the remaining trials the fixation dot remained dark gray. See the description of tasks below for how the colors were controlled.

Each character string was presented exactly once to each participant during the experiment. The sets of words used for each participant were also balanced to equalize the orthographic neighborhood size, log lexical frequency, mean reaction time and accuracy estimated from the English Lexicon Project (Balota et al., 2007) across all conditions in the experiment. We generated 6 sets of word-to-location assignments, and randomly used one for each participant.

#### Task

Participants performed three different tasks during different runs, two of which required attending to the character strings, and one that encouraged participants to ignore them. In the ***lexical decision task,*** participants reported whether the character string on each trial was a real word or not. In the ***stimulus color task***, participants reported whether the color of the character string was red or gray. In the ***fixation color task***, participants reported whether or not the fixation dot turned red.

The fixation color task served as the baseline for neural responses: the same stimuli were presented as in the other two tasks, but the participants were required to ignore them as they tried to detect a subtle, threshold-level color change that occurred at the exact same time the character string appeared. The 150 ms duration did not allow for attention to switch from the fixation dot to the character string. For the stimulus color task and the fixation color task, the saturation of the red color was updated using an adaptive staircase that converged at the 80% correct detection threshold. During runs when the character (or fixation dot) color was not task-relevant, the saturation levels across trials were replayed from the staircase in another color task run.

Across all three tasks, the probability of a real word appearing was 0.33. The probability that the character string was red was 0.33. Similarly, the probability that the fixation dot turned red was 0.33. All three of those events were independent of each other. Thus, for all three tasks, the probability of a “target” (a real word, a red character string, or a red fixation dot) was 0.33. All that changed across runs was which stimulus attributes were task-relevant.

Participants responded via a button press with their right index finger for “yes” responses (word or not, red or not), and right middle finger for “no” responses. Participants were instructed to respond within 3s after each stimulus and to prioritize accuracy over speed.

#### Procedure

Before the first scanning session, participants were trained on all tasks in a testing room outside the scanner. They performed a minimum of 24 trials of each task, and were also trained to calibrate the eye-tracker and to maintain fixation. Instructions reminded them to respond as accurately as possible.

The main experiment consisted of fifteen 5-minute runs, with 5 runs for each of the three tasks. The order in which the tasks were presented was:

LD SC FC SC FC LD FC LD SC LD FC SC SC LD FC

where LD is lexical decision, SC is stimulus color, and FC is fixation color task. This order was chosen such that each task was equally likely to follow both of the others.

Each run began with written instructions on the screen, informing the participant which task to perform. They pushed a button to proceed, and then were quizzed on which task they were to perform in that run. The run would not start until the participant answered correctly, to ensure that they performed the correct task.

There was no feedback about response accuracy after each trial, but at the end of each run, text appeared on the screen to report their percent correct, along with the percentage of trials in which they failed to make a response.

The sequence of trials within each run was counterbalanced and uniquely generated for each participant. There were 45 trials in each run, and the category of the character string stimulus was randomized. In summary, there were 27 unique conditions, defined by stimulus type, stimulus location, and task. We collected 25 trials per condition.

#### Predictions

If the VWFA is sensitive only to stimulus features, it should respond more strongly to words than false fonts by the same degree across tasks. If it is sensitive only to linguistic task demands, it should respond most strongly during the lexical task regardless of stimulus type. Other hypotheses involve interactions; for instance, the VWFA’s preference for words may be due to top-down feedback and arise only when the stimuli are attended. It is difficult to predict whether such modulations would occur only during an explicitly linguistic task, or whether they would occur *automatically* whenever the stimuli are attended (as in the stimulus color task). Moreover, the response to false fonts may even be suppressed when attended, similar to what was found during a shape judgement task (White et al., 2023).

### Localizer for visual category-selective ventral temporal regions

We examined activity in regions of interest (ROIs) that we defined precisely in each individual’s brain using independent localizer scans (**Figure 1A**). This is necessary because the VWFA is composed of several small patches that differ across individuals (Centanni et al., 2019; Caffarra et al., 2021). Averaging in a template brain obscures word-selective processing (Glezer and Riesenhuber, 2013; Mahowald and Fedorenko, 2016; Stevens et al., 2017). An example participant’s ROIs are shown in **Figure 1B**.

#### Stimuli

Participants viewed sequences of images, each of which contained 3 items of one category: words, pseudowords, false fonts, faces, and limbs (examples in **Figure 1A**). This localizer was also used in White et al., 2023, which describes the stimuli in detail. The major change here is the addition of the “limbs” category instead of objects. The grayscale limb images included hands, arms, feet and legs (Stigliani et al., 2015).

#### Task and Procedure

Participants performed a one-back repetition detection task. On 33% of the trials, the exact same images flashed twice in a row. The participant’s task was to push a button with their right index finger whenever they detected such a repetition. Each participant performed 4 runs of the localizer task. Each run consisted of 77 four-second trials, lasting for approximately 6 minutes. Each category was presented 56 times through the course of the experiment.

### Language Network Localizer

#### Stimuli

In order to map the canonical language network, we used a language localizer (Mahowald and Fedorenko, 2016). Details about the localizer are provided by the authors who developed the resource. Briefly, the stimuli on each trial were a sequence of 12 written words or pronounceable pseudowords, presented one at a time. The words were presented as meaningful sentences, while pseudowords formed “Jabberwocky” phrases that served as a control condition.

#### Task and Procedure

Participants were instructed to read the stimuli silently to themselves, and also to push a button upon seeing the icon of a hand that appeared between trials. Participants performed three runs of the language localizer. Each run included 16 trials and lasted for 6 minutes. Each trial lasted for 6s, beginning with a blank screen for 100ms, followed by the presentation of 12 words or pseudowords for 450ms each (5400s total), followed by a response prompt for 400ms and a final blank screen for 100ms. Each run also included 5 blank trials (6 seconds each).

### Study Procedure

The entire study required four sessions per participant. The first session was dedicated to obtaining informed consent, MR safety screening and training the participant on the tasks and eye-tracking. The first two MRI sessions were identical: T1 structural scan, 6 runs of the main experiment, 2 runs of the ventral visual localizer, and finally fieldmap acquisition. One run of the localizer was interspersed after every 2 runs of the main experiment. In the third session, participants performed the final 3 runs of the main experiment, and 3 runs of the language localizer.

### Analysis of behavioral performance

We calculate accuracy as the proportion of trials with correct responses made within 3.85 seconds after stimulus onset. On an average of 8% of trials (SEM=2%), the participant did not make a response within that window. Those trials were excluded from analyses of accuracy and response time.

### Preprocessing

We pre-processed our MRI data with *fMRIPrep* 21.0.1 (Esteban et al., 2019), which is based on *Nipype* 1.6.1 (Gorgolewski et al., 2011).

#### Structural data preprocessing

For each subject, for each session, we obtained a T1-weighted structural scan. These images were skull-stripped, corrected for intensity non-uniformity and averaged across sessions. Cortical surfaces were constructed from the boundaries between gray and white matter using Freesurfer’s implementation (Dale et al., 1999; Reuter et al., 2010). We obtained surfaces in subjects’ anatomical space as well as the template *fsaverage* space.

#### Functional data preprocessing

Head-motion parameters with respect to the BOLD reference (transformation matrices, and six corresponding rotation and translation parameters) were estimated before any spatiotemporal filtering using mcflirt (FSL 6.0.5.1:57b01774, Jenkinson et al. 2002). Each functional EPI sequence acquired during a single session was corrected for magnetic field inhomogeneities using a B_0_ nonuniformity map collected during the same session. All runs were aligned to native anatomical (*fsnative*) and template MNI and *fsaverage* spaces using Freesurfer. The runs were slice-time corrected (using 3dTShift from AFNI) and then co-registered to the T1-weighted images (using bbregister from Freesurfer) (Greve and Fischl, 2009). The BOLD time-series were also resampled onto each subject’s native space cortical surface (Lanczos, 1964).

### BOLD Response Estimation

To estimate the BOLD responses to each trial, we used GLMSingle in Python (Prince et al., 2022) which optimizes the hemodynamic response function for every voxel or node in the brain. For the main experiment, the design matrix consisted of a column for each combination of task, stimulus and presentation location. For the functional localizer, each unique category of stimulus received its own column. For the language localizer, the design matrix included two columns corresponding to the sentence and jabberwocky conditions. Before estimating the BOLD responses, the timeseries for each run were upsampled from 1.5s to 1s using an interpolation function from pyslicetime (Kay et al., 2020). We used GLMSingle to estimate the response to each individual trial of each condition for each surface node in each participant’s fsnative space. The algorithm found the optimal hemodynamic responses function for each surface node, estimated cross-validated nuisance regressors computed from repetitions of the same experimental condition, and regularized responses for each node to remove correlated noise across single trials.

### Region of Interest Definition

All regions of interest (ROIs) analyzed in this study were obtained using one of three procedures: (1) The ventral visual localizer scan, used to define the regions depicted in color in **Figure 1B**; (2) The language localizer scan, to define regions outlined in black; or (3) from an independent study using a separate pool of subjects (White et al., 2023), for the regions outlined in white.

We defined visual category-selective ROIs in the ventral temporal cortex by contrasting the responses to different image categories. For each category (words, faces, and limbs), we computed t-statistics for the contrast of that category vs. all others using a paired-sample t-test. The t-values were visualized on each participant’s inflated cortical surfaces, with the threshold at t ≥ 3.

We identified up to three text-selection areas (the VWFAs) in each hemisphere. VWFA-1 and VWFA-2 were in the occipitotemporal sulcus, VWFA-1 being anterior to hV4 and VWFA-2 as a separate region anterior to VWFA-1. The face-selective areas, FFA-1 and FFA-2, were medial to the VWFAs, within the fusiform gyrus. FFA-2 was anterior to FFA-1 but sometimes contiguous with it. The limb-selective area was lateral to the VWFAs, defined in the occipito-temporal sulcus (Grill-Spector and Weiner, 2014) but in some subjects contiguous with a patch on the lateral side of the brain (encompassing what is sometimes called the extrastriate body area (Downing et al., 2001)). The ROI for area hV4 was taken from a previous study that conducted retinotopic mapping (White et al., 2023), defined in fsaverage space and remapped into each individual’s native surface.

We also defined two language-responsive regions in the left frontal lobe (“Broca’s area”). First, using our data from the language localizer, we used a similar procedure as the one described above, with a contrast between sentence and jabberwocky conditions. For each individual, we defined a region with a peak in the inferior frontal sulcus, extending into the inferior frontal gyrus as the “Language Broca’s Area” (Fedorenko et al., 2010). This is the black frontal region in **Figure 1B**. The second left frontal area we took from a previous study that contrasted responses to written words vs. strings of shapes (White et al., 2023). This area was defined in fsaverage space and remapped into each individual’s native surface. We call “Word Form Broca’s,” and it is the white frontal region in **Figure 1B**. It was in the left precentral sulcus, slightly posterior to “Language Broca’s. The names of these two frontal language-related regions are intended to reflected how these ROIs were defined, and not to make any strong claims about their function. Given the variety of neuroimaging tasks the frontal operculum is active in, it is unlikely that these regions serve any singular processing function. Finally, we also leveraged the language localizer to define an ROI in the superior temporal sulcus (STS).

ROIs on an example subject’s cortical surfaces are shown in **Figure 1B**, and all participants are shown in **Extended Data Figure 1-2**. Details about the numbers of participants in which we were able to identify each ROI are reported in **Table 1**.

**Table 1:**
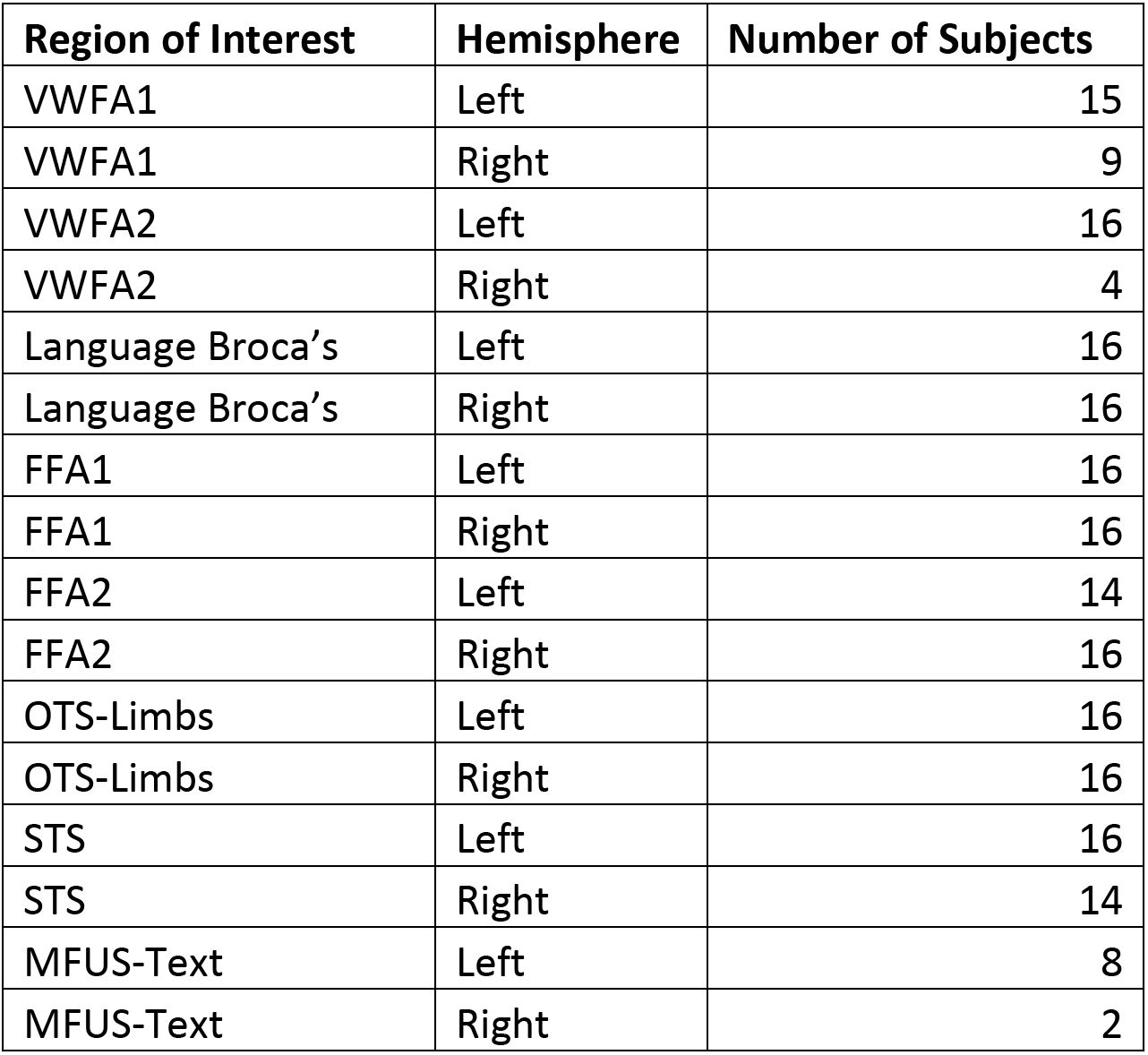
Number of participants with each ROIs: Number of participants (N) in whom each ROI could be defined, reported separately for each hemisphere. Two other ROIs (V4 and Word Form Broca’s) were defined using data from White et al. 2023 and applied to all participants.

To create the group-level ROIs in **Figures 5**-**7**, we used the following approach. For the VWFAs, we projected individually-defined contrasts between responses to text vs. all other categories in the visual localizer to the template fsaverage space. We then concatenated these contrasts across all subjects, and performed a one sample t-test across subjects to localize the nodes in which responses between text and other categories were significantly different from zero. We performed an FDR correction for multiple comparisons, and saved the nodes with significant t-values. We used this map to define labels for VWFA-1 and -2 in the fsaverage space. For the Language Broca’s area, we used a similar approach but with the contrast between meaningful sentences and jabberwocky in the language localizer.

### Statistical Analysis

We used linear mixed effect models to determine which of our experimental manipulations significantly affected BOLD responses. The dependent variable was the mean beta response for each subject, and the fixed effects included the stimulus type, location and task. The random effects included slopes and intercepts for these variables across participants. The F-statistics and p-values reported were obtained with a type-three sum of squares table upon performing repeated-measures ANOVA using orthogonalized contrasts on the fixed effects. The t-tests for post-hoc tests are reported with FDR corrected p-values. All t-tests conducted were two-tailed. These analyses were implemented using the python package pymer4 (Jolly, 2018). We also report Bayes Factors (BFs) for each pairwise test, to quantify strength of evidence. The BF is the ratio of the probability of the data under the alternate hypothesis (that two conditions differ), relative to the probability of the data under the null hypothesis (Rouder et al., 2009). For example, a BF of 10 indicates that the data are ten times more likely under the alternate hypothesis than the null hypothesis. We computed BFs using the R package “BayesFactor” (Morey et al., 2015).

### Whole-cortex Analyses

We performed several analyses of cortex-wide activity to complement our approach of individually-mapped ROIs. The goals were to create maps of the sensitivity to stimulus types or task demands. For each analysis, we first obtained the test statistic of interest for each participant in their native anatomical space (e.g., the mean difference in BOLD response magnitude for text vs. false fonts in a particular task). Next, we transformed this map of statistics to the *fsaverage* template. Finally, for each cortical surface node in *fsaverage*, we conducted a one sample t-test over subjects, to find which nodes had a value different from zero. We then performed FDR correction on these p-values, and thresholded the resulting map at p<0.05. This was all done with custom Python code, using scipy (Virtanen et al., 2020).

### Functional Connectivity Analysis

The goal of this analysis was to quantify the correlations between brain regions in terms of the *fluctuations* of BOLD response across trials, separately for each experimental condition. Similar to previous studies (Kay and Yeatman, 2017; White et al., 2023), we evaluated across-region correlations in the single-trial beta responses, from which low frequency drifts and other nuisance regressors had already been removed, rather than the raw time series. To do so, for each participant, for each surface node or ROI, we collated vectors of the responses to trials of each condition. Conditions are defined by the combination of task, stimulus type, and stimulus location. (Trials with real words and pseudowords were treated separately). For each condition, we subtracted out the mean response magnitude from the vector of responses, yielding de-meaned “residuals”.

We then analyzed across-region correlations in these residual responses in two ways. First, we conducted whole-brain analyses of functional connectivity to a “seed region,” in each individual’s native cortical surface. We first averaged responses over the nodes within the seed region, and then we calculated the Pearson correlation coefficient between the seed’s residuals and the residuals from every other surface node. We then projected the resulting correlation coefficients from each subject’s native space to the *fsaverage* space. With all 16 participants’ correlation maps in the same space, we averaged the correlation coefficients and at each node computed a one-sample t-test to assess whether the correlation was significantly different from Using the FDR-corrected p-values for each vertex as a threshold, we obtained a map of nodes that were significantly correlated with the seed region during each condition. We performed these analyses with the scipy and statsmodels libraries (Virtanen et al., 2020).

Second, we assessed how functional connectivity between pairs of ROIs changed across experimental conditions. In **Figure 6B**, we focused on the correlated activity between the VWFAs and the “Language Broca’s area”, given the result of the whole-brain analysis and prior evidence (White et al., 2023). The correlations between residuals were calculated on data first averaged across nodes within the two ROIs in each individual’s native brain, separately for each condition. We then averaged the correlation coefficients across subjects. For these measures we conducted similar statistical tests as for the BOLD response magnitudes (described above).

## Results

### Behavior

The participant’s task on each trial was to push one of two buttons to report whether or not they saw a ‘target’: a real word in the lexical decision task; red letters in the stimulus color task; or a brief change in the fixation dot’s color from gray to red. In both color tasks, an adaptive staircase continuously adjusted the saturation of red color to each participant’s detection threshold. The character string’s position varied randomly across trials, either centered on the fixation mark, 3° left, or 3° right. The stimulus types were randomly intermingled, and the task conditions were blocked.

**Figure 1D** plots mean task accuracy in each condition. Using linear mixed models with subjects as random effects, we analyzed accuracy as a function of the task, stimulus type, and stimulus location. We found a main effect of all three experimental manipulations (Task: F(2,33)=37.77, p<0.001, Stimulus Type: F(2,23)=109.84, p<0.001, Stimulus Location: F(2,46)=6.12, p=0.004). There were also and two-way interactions between task and stimulus type (F(4,375)=117.09, p<0.001), and between task and location (F(4,375)=9.8, p<0.001). Accuracy was highest in the stimulus color task, and lowest in the fixation color task (all pairwise comparisons p<0.03). In the lexical decision and stimulus color tasks, accuracy was higher when the stimuli were presented at fixation (0°) than ±3° in the parafovea, but the opposite was true in the fixation task. In other words, fixation task accuracy relatively impaired by characters appearing at fixation (**Figure 1D**). Post-hoc tests (corrected for false discovery rate) revealed better lexical decision performance when real words were presented in the right compared to the left hemifield (t=2.31, p=0.03), consistent with prior results. Reaction times are plotted in **Figure 7**.

### The VWFA prefers text over unfamiliar characters even when attention is withdrawn

The mean BOLD responses in the left VWFA are shown in **Figure 2**. In all our analyses, we found that the two main subregions of the left VWFA (VWFA-1 and VWFA-2) had very similar response patterns. We therefore average across them here. Results for the separate regions are reported in **Extended Data Table 2-1.** A third ventral text-selective region, MFUS-text, is summarized in **Figure 3F**.

**Figure 2:**
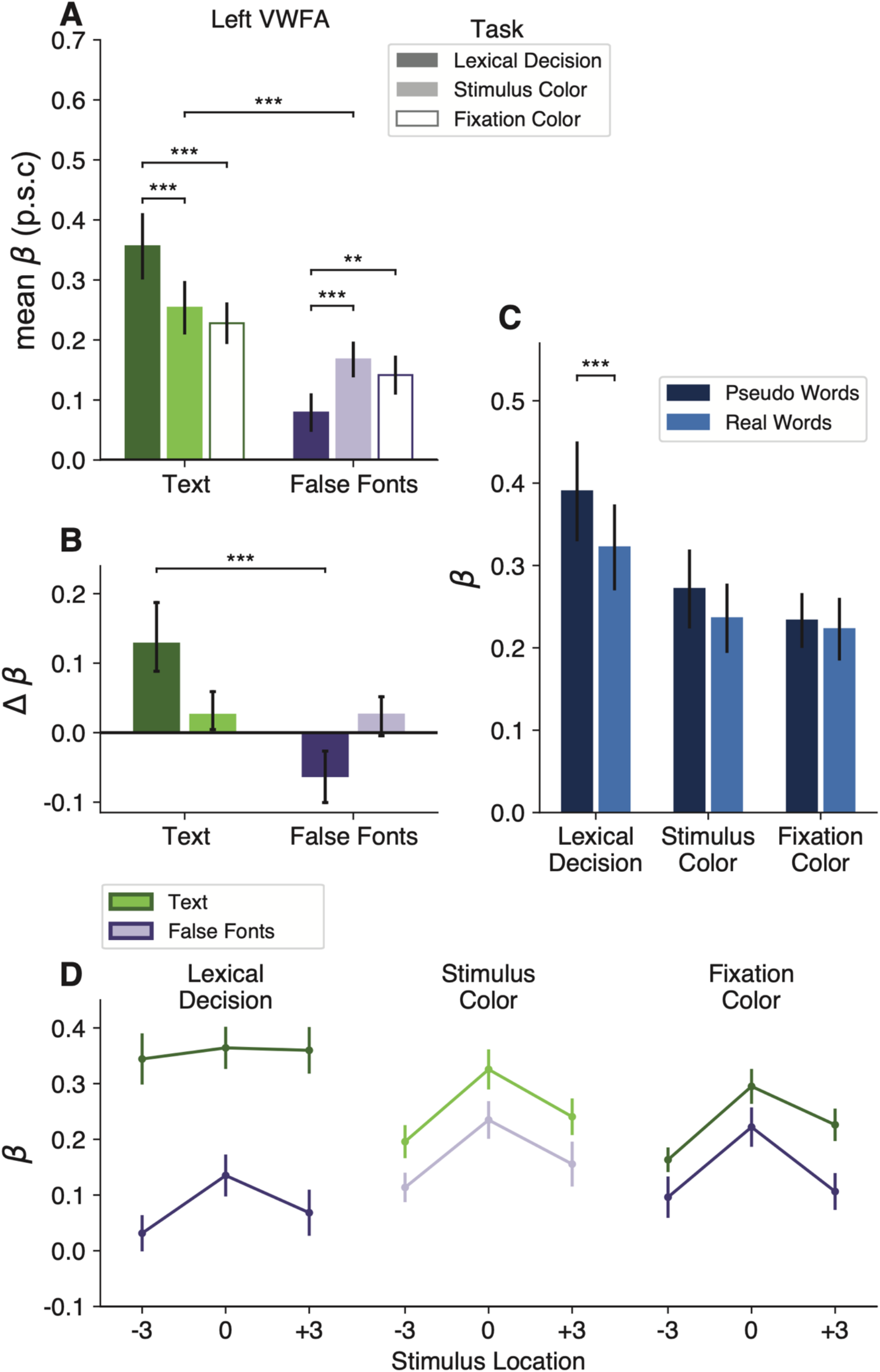
The VWFA’s activity reflects an interaction of stimulus features and task demands. **(A)** Mean percent signal change (p.s.c.) as a function of stimulus type and task condition in left hemisphere VWFAs. “Text” includes both real words and pseudowords (which are separately analyzed in panel **C**). Error bars represent ±1 SEM. Horizontal bars and stars indicate statistical significance for the main effect of stimuli and pairwise t-tests between tasks. **(B)** Task effects computed as the mean difference in percent signal change from the fixation task as baseline. Error bars indicate *95% confidence intervals.* Horizontal bars and stars indicate a significant difference in the effect of engaging in the lexical task (compared to fixation task) on responses to text vs. false fonts. The effect of the stimulus color task did not differ across stimulus types. For task differences in individual participants, see Extended Data Figure 2-1. **(C)** Responses to words vs. pseudowords in the three tasks. The lexicality effect is only observable during the lexical decision task. Error bars are ±1 SEM. **(D)** Responses in the VWFAs as a function of stimulus position, separately for each task and stimulus type (green = text; purple = false fonts). Error bars are ±1 SEM.

**Figure 3:**
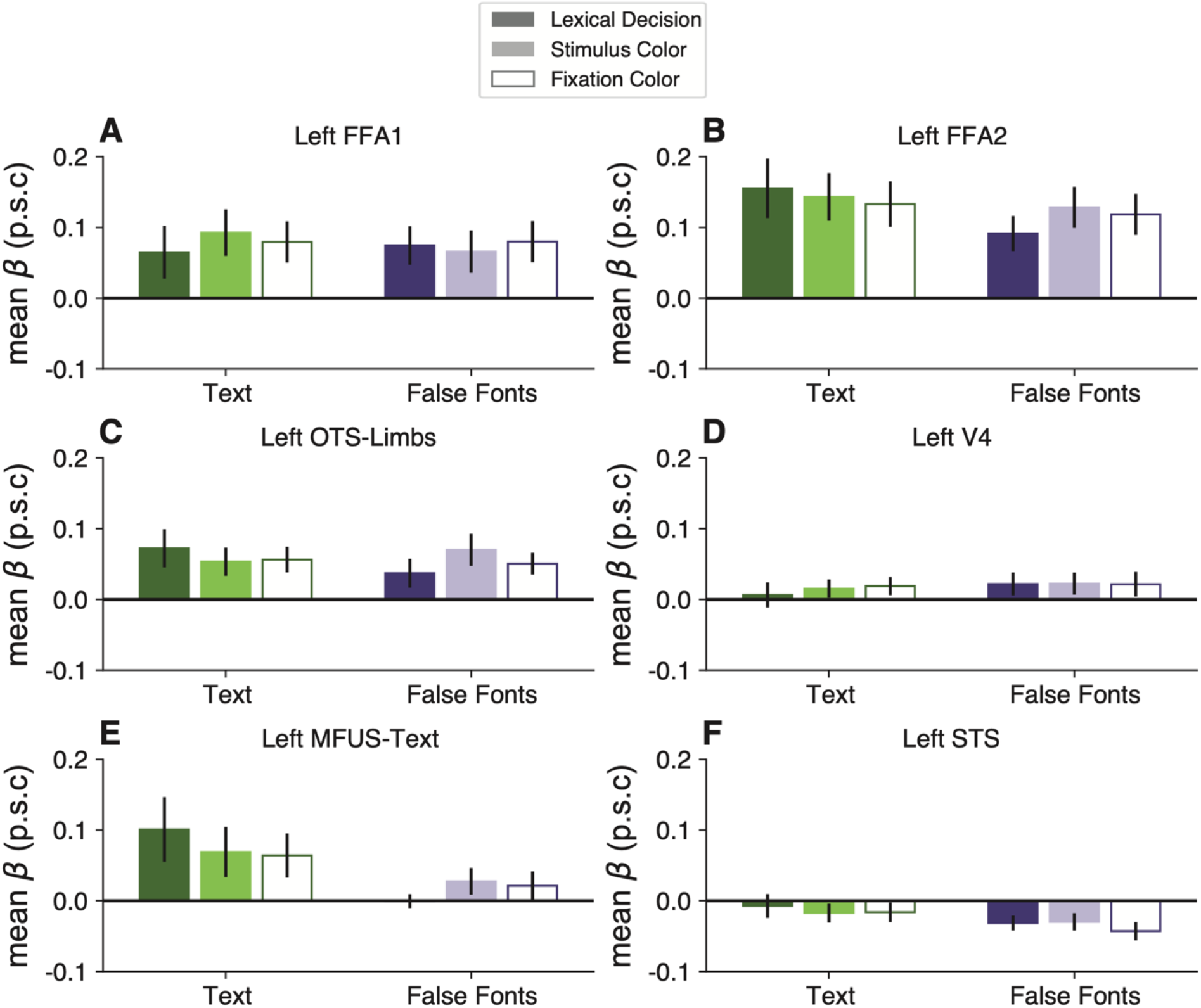
Task and stimulus effects are absent in other regions of left occipital and temporal lobes. Each panel plots mean BOLD responses in one ROI, formatted as in Figure 2A. See Extended Data Figure 3-1 for statistics on these control regions and Extended Data Figure 3-2 for right hemisphere ROIs.

**Table 2:**
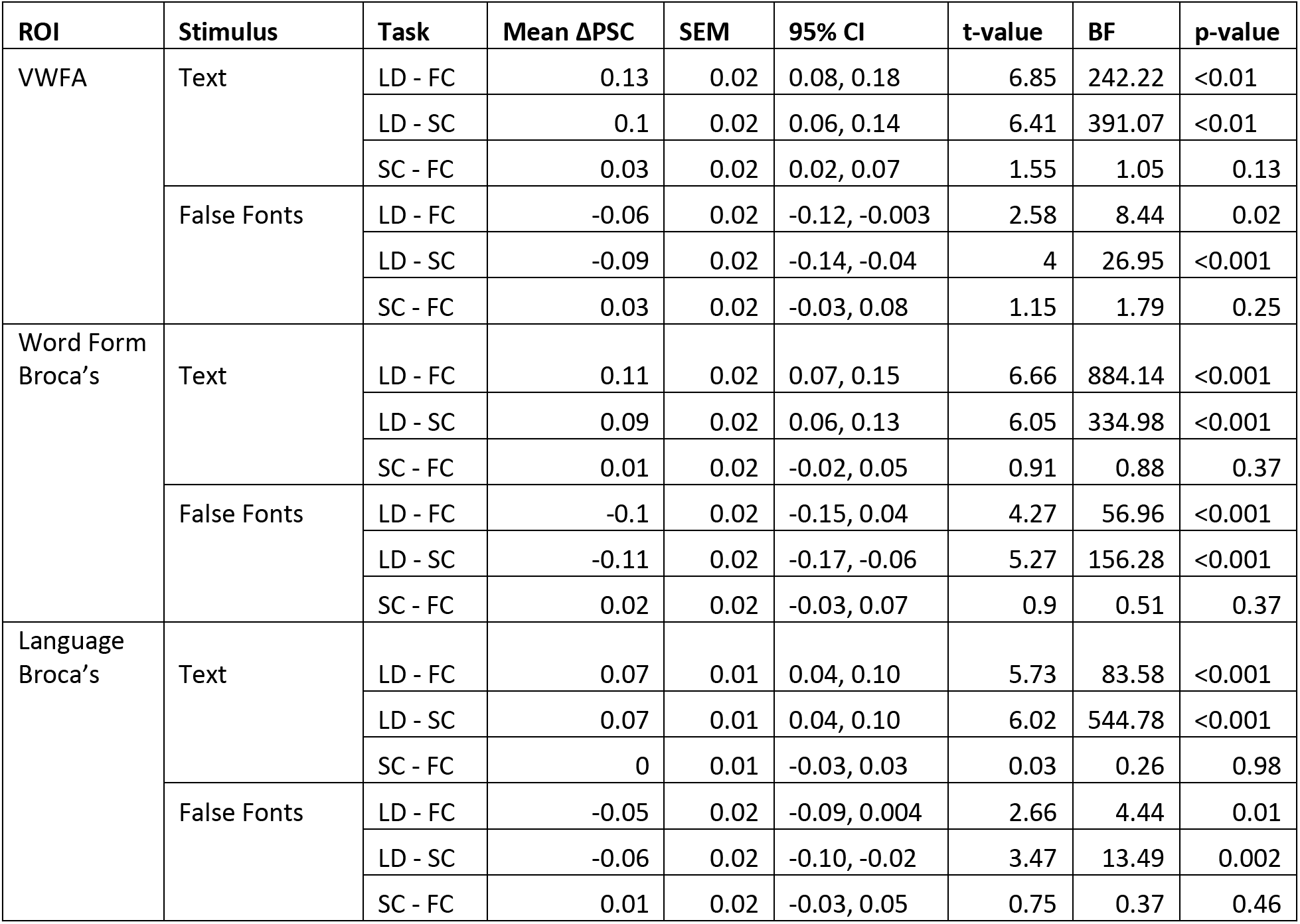
Mean task effects on percent signal change: Across pairs of tasks, separately for text and false font stimuli, in the left VWFA and the two Broca’s areas. See Extended Data Table 2-1 for statistics on VWFA-1 and VWFA-2 separately. For each ROI, the six p-values are corrected for false discovery rate. Abbreviations: PSC = Percent Signal Change, SEM = Standard Error of Mean, CI = Confidence Interval, LD = lexical decision task; FC = fixation color task; SC = stimulus color task; BF = Bayes Factor.

The first result concerns the bottom-up selectivity for text in the VWFA. This region was defined as a patch of cortex that, in an independent localizer scan, responded more strongly to attended words than to false fonts, faces, and limbs (during a one-back task). We therefore expected that the VWFA would also prefer words and pseudowords over false font strings in the main experiment. But other researchers have argued that this region appears to have selectivity for text due to top-down influence from cerebral networks involved in language and attention (e.g., Dȩbska et al 2023; Price & Devlin, 2011). Such an account predicts that during a non-linguistic task in which attention is totally withdrawn from the stimuli, the VWFA would respond equivalently to words as to meaningless shapes that have similar visual features.

To test that prediction, we compared the response to text (words and pseudowords together) vs. false font strings during the fixation color task. In that task, the stimuli were task-irrelevant and the participant was focused on detecting a threshold-level color change in the fixation dot. They did not have time to switch attention from the dot to the character strings that flashed briefly at the same time. Nonetheless, the left VWFAs responded more strongly to text than false fonts during this fixation task (mean betas = 0.23 vs 0.14; t(15)=4.82, p = 0.0002, BF=139.87). These data are the two open bars in **Figure 2A**. This text selectivity was maintained even when the stimuli were presented 3° from the focus of attention at the fixation dot, as shown in the bottom right panel of **Figure 2D**

We infer that the VWFA has inherent selectivity for a certain category of shapes: strings of familiar letters. That selectivity is minimally dependent on attentional engagement with the stimuli or on voluntary linguistic processing. Importantly, however, the selectivity for text was absent in the FFAs, limb-selective areas, and V4 (see **Figure 3**).

### The lexical decision task enhances the VWFA’s response to text and suppresses its response to false fonts

The next result concerns how the VWFA’s activity changed when the participant engaged in the lexical decision task, which required them to attend to each stimulus and judge whether it was a known English word. To start, we found a significant interaction between stimulus type and task on mean BOLD response magnitudes (F(2, 108)=34.34, p=3x10^-12^). As shown in **Figure 2A-B**, the effects of the task differ across stimulus types. To quantify that interaction, for each stimulus type we calculated comparisons of the responses between pairs of tasks. The statistics are reported in **Table 2**. The mean differences from the fixation task baseline are shown in **Figure 2B**, with error bars that represent 95% confidence intervals.

First, compared to when the stimuli were ignored during the fixation color task, the VWFA’s response to text (real words and pseudowords) was enhanced during the lexical decision task, but not during the stimulus color task. The dark green bar in **Figure 2B** shows the enhancement in the lexical task. The mean response during the lexical decision task was 1.6 times the mean response during the fixation color task. All individual participants showed this effect.

Second, for false fonts, we observed the opposite task effect: responses during the lexical decision task were *lower* compared to the other two tasks. See the dark purple bar in **Figure 2B**. This result suggests that VWFA’s response to non-preferred stimuli is suppressed below the sensory baseline level while the brain is engaged in linguistic processing. 12 out of 16 participants showed lower responses for false fonts during the lexical decision task, compared to the fixation color task baseline. The mean response to false fonts in the lexical task was 54% of the mean response to false fonts in the fixation task. To characterize the consistency of these effects, we refer the reader to **Extended Data Figure 2-1**, which compares the VWFA responses for each task and stimulus type within individual participants.

Moreover, **Figure 2C** shows that the VWFA responds differently to real words and pseudowords, but only during the lexical decision task (word type by task interaction: F(2,45)=3.5, p=0.038). Consistent with prior studies, pseudowords evoked larger responses than real words during the lexical decision task (t(15)=2.8, p = 0.01, BF=4.58). That may be because processing pseudowords takes more time than processing familiar words (Vinckier et al., 2007; Cohen et al., 2008). However, that *lexicality effect* is absent when the stimuli are ignored during the fixation task (t(15)=0.6, p = 0.56, BF=0.3). This result suggests a key role for top-down input in creating the VWFA’s sensitivity to higher-level lexical features of the stimuli (in contrast to the selectivity for letter strings, which was strong in all tasks).

**Figure 2D** shows that VWFAs are also sensitive to the visual field position of the words (Rauschecker et al., 2012; Le et al., 2017; White et al., 2023). In general, the VWFA prefers stimuli at the fovea (0°) over stimuli presented 3° to the left or right (main effect of location: F(2,24)=29.7, p < 0.001). However, that spatial sensitivity is greatly reduced for text presented during the lexical decision task. In **Figure 2D**, compare the dark green line on the left to the lighter green lines on the right. During the lexical task, there were no significant differences in pairwise comparisons of stimulus positions (all t<1, p>0.7), because the enhancement of attended words was strongest at ±3°. Thus, voluntary effort to read enhances the response especially when words are in the parafovea where legibility is reduced (Vinckier et al., 2007; Kay and Yeatman, 2017). Differences between the lexical decision and fixation color tasks were also present when the stimuli were presented foveally, for text (t(15)=3.75, p=0.0007, BF=43.76) as well as for false fonts (t(15)=2.69, p=0.02, BF=3.59).

### Top-down modulations do not occur automatically when words are visually attended

Word recognition is sometimes described as an “automatic” process (Stroop, 1935; Price et al., 1996; Brown et al., 2002; Ruthruff et al., 2008; Pattamadilok et al., 2017; Joo et al., 2021). The Stroop effect is the most famous evidence: participants cannot stop themselves from processing the meaning of a word when instructed to say aloud the color of its ink. An automaticity-based account would predict that attending to words to judge the font color would engage the VWFA just as strongly as when voluntarily reading the words. However, we found that the VWFA responded quite similarly during the stimulus color task as when the words were *ignored* during the fixation task. The light bars in **Figure 2B** represent the mean differences across those two tasks for text (green bar) and false fonts (purple bar). While engaging in the lexical decision task greatly increased the response to text, engaging in the stimulus color task had a negligible effect (not significant after correcting for multiple comparisons; BF=1.05). The response to false fonts was not suppressed in the stimulus color task, but rather slightly and not significantly increased (BF=1.79; see **Table 2**).

One potential criticism of the comparison between the lexical and stimulus color tasks is that the latter could be accomplished by attending to only one letter. However, the character string was presented for only 150 ms at an unpredictable location, and with the saturation of the red color set to detection threshold. The optimal strategy therefore was to integrate the color signal across all the letters.

Thus, the elevation of the VWFA’s response to words does not occur automatically when the words are attended, but requires engagement in an explicitly lexical task. The same applies to the relative suppression of the response false fonts.

### The task and stimulus interactions are absent from other occipital and temporal regions

We analyzed responses in six other regions of interest in the left occipital and temporal lobes. See **Figure 3**. Within the ventral occipito-temporal cortex, we analyzed V4, face-selective regions in the posterior and middle fusiform gyrus (FFA-1 and FFA-2, respectively), a limb-selective region (OTS-Limbs), and another text-selective region within the mid-fusiform gyrus (MFUS-Text, which was only detectable in half of our participants). Additionally, we analyzed responses in a large swath of the superior temporal sulcus (STS) that was activated in the language localizer.

We did not find a significant effect of task, stimulus type, or an interaction between task and stimuli within any of these ROIs. Thus, the pattern of activity we found in the VWFA is unique among the visual regions we analyzed and also absent in the STS. Statistics for all ROIs are reported in the **Extended Data Figure 3-1**. FFA-2 had a pattern that was qualitatively similar to the VWFA, but the task effects were less consistent across subjects and not statistically significant (all *p*s>0.5).

The only marginally significant effect was observed in the left MFUS-Text (**Figure 3F**), where responses to text were somewhat higher in the lexical task than the other two tasks (p=0.07). Left MFUS-text also responded more strongly to text than false fonts overall, but not significantly after correcting p-values across the ROIs, perhaps because we were able to localize this region in only 8/16 participants. Right hemisphere regions, including VWFA, are shown in **Extended Data Figure 3-2**. There were no significant effects of stimulus or task in any of these regions.

### Broca’s area shows strong task-dependent enhancement and suppression

We also localized two language processing regions in the left frontal cortex. The first we refer to as “*Language Broca’s*,” because it was defined in each individual by an independent language localizer (see Methods) (Fedorenko et al., 2010; Mahowald and Fedorenko, 2016). It was located in the inferior frontal sulcus, sometimes extending into the inferior frontal gyrus. The second region we refer to as “*Word Form Broca’s*” because it was defined (in an average brain) from a previous study contrasting the response to written words and non-letter shape strings (White et al., 2023). It was slightly posterior to Language Broca’s, in the precentral sulcus. A similar contrast on our main experiment, contrasting text stimuli with false fonts, produced an overlapping region. We used the Word Form Broca’s from the previous study to avoid using the same data for defining an ROI and analyzing mean responses. The locations of the two Broca’s regions relative to each other are depicted in **Figure 1B** in an example subject, and on the average brain in **Figure 5B**.

**Figure 4** shows that in both these Broca’s area sub-regions, we found interactions between stimulus type and task that went in the same direction as in the VWFAs. However, response magnitudes in the more anterior “Language Broca’s area” were different: it did not respond positively to stimuli overall (**Figure 4A**). Its response differed from baseline only during the lexical decision task, but in opposite directions for text vs. false fonts: a strong increase for text, and a decrease below baseline for false fonts (**Figure 4C**). These differences between the lexical decision task and the fixation color task were observed for text in 15 out of 16 participants, and for false fonts in 13 out of 16 participants. It also showed sensitivity to lexicality (pseudowords > real words) only during the lexical task (**Figure 4E**).

**Figure 4:**
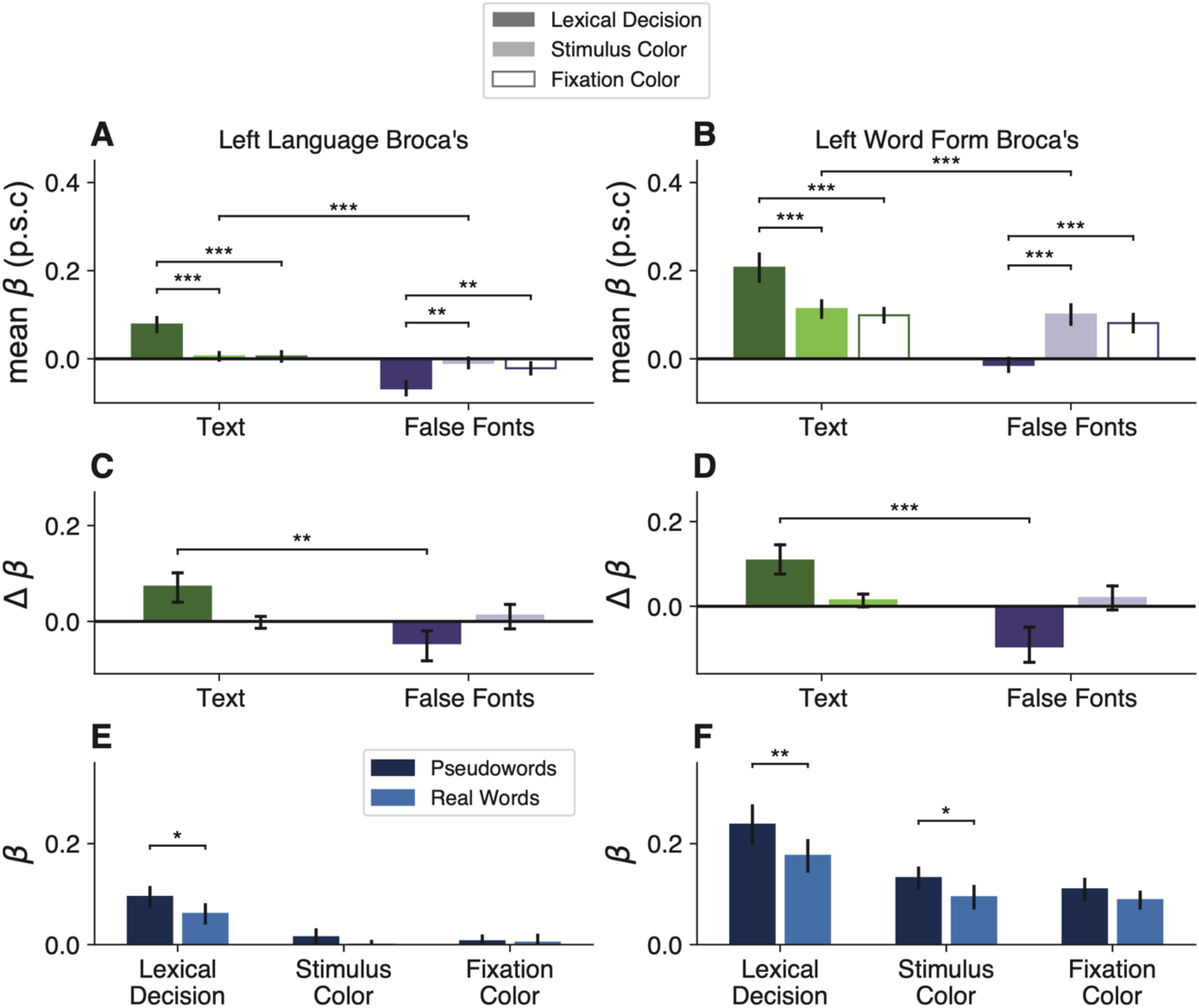
Task and stimulus effects in two left frontal regions. (**A, B**) Mean percent signal change as a function of stimulus type and task in two frontal regions in the left hemisphere. Format as in Figure 2A. **(C, D)** Difference in mean percent signal change from fixation task baseline. Format as in Figure 2C; error bars indicate *95% confidence intervals.* **(E, F)** Responses to real words vs. pseudowords in each task (format as in Fig 2E).

The response magnitudes in Word Form Broca’s were more similar to the VWFA, with generally positive trial-evoked responses above baseline (**Figure 4B**). It showed the same task effects: a selective enhancement for text and suppression for false fonts during the lexical decision task (**Figure 4D**). In this region, we observed the task effects for text in all participants, and for false fonts in 14 out of 16 participants. It also responded more strongly to pseudowords, except during the fixation task (**Figure 4F**). Statistics for the task effects in these two regions are reported in **Table 2**.

One interpretation of these data is that the “Language Broca’s area” functions as a *control region* that applies positive or negative modulations to other parts of the reading network. This region is engaged only during the lexical task. If letter strings are presented, its activity rises and it in turn excites other regions involved recognizing words: the VWFAs as well as the more posterior Word Form Broca’s. If it detects false fonts during the lexical task, its activity decreases below baseline and the other regions are suppressed as well.

### A whole-cortex analysis of the task-by-stimulus interaction highlights Broca’s area and the left VWFAs

The preceding analyses were based on small regions of interest localized precisely in each individual’s brain from independent data. This was necessary given the documented shortcomings of first morphing to an average brain. However, we also investigated whether our primary findings are widespread across the cortex or in fact unique to the primary regions of interest.

A key result in the VWFA and Broca’s ROIs was an interaction between stimulus type and task: the response to text was stronger in the lexical task than in the fixation color task, but the response to false fonts was stronger in the fixation task. **Figure 5** shows a whole-cortex map of this interaction between stimulus type (text vs false fonts) and task (lexical decision vs fixation task).

**Figure 5:**
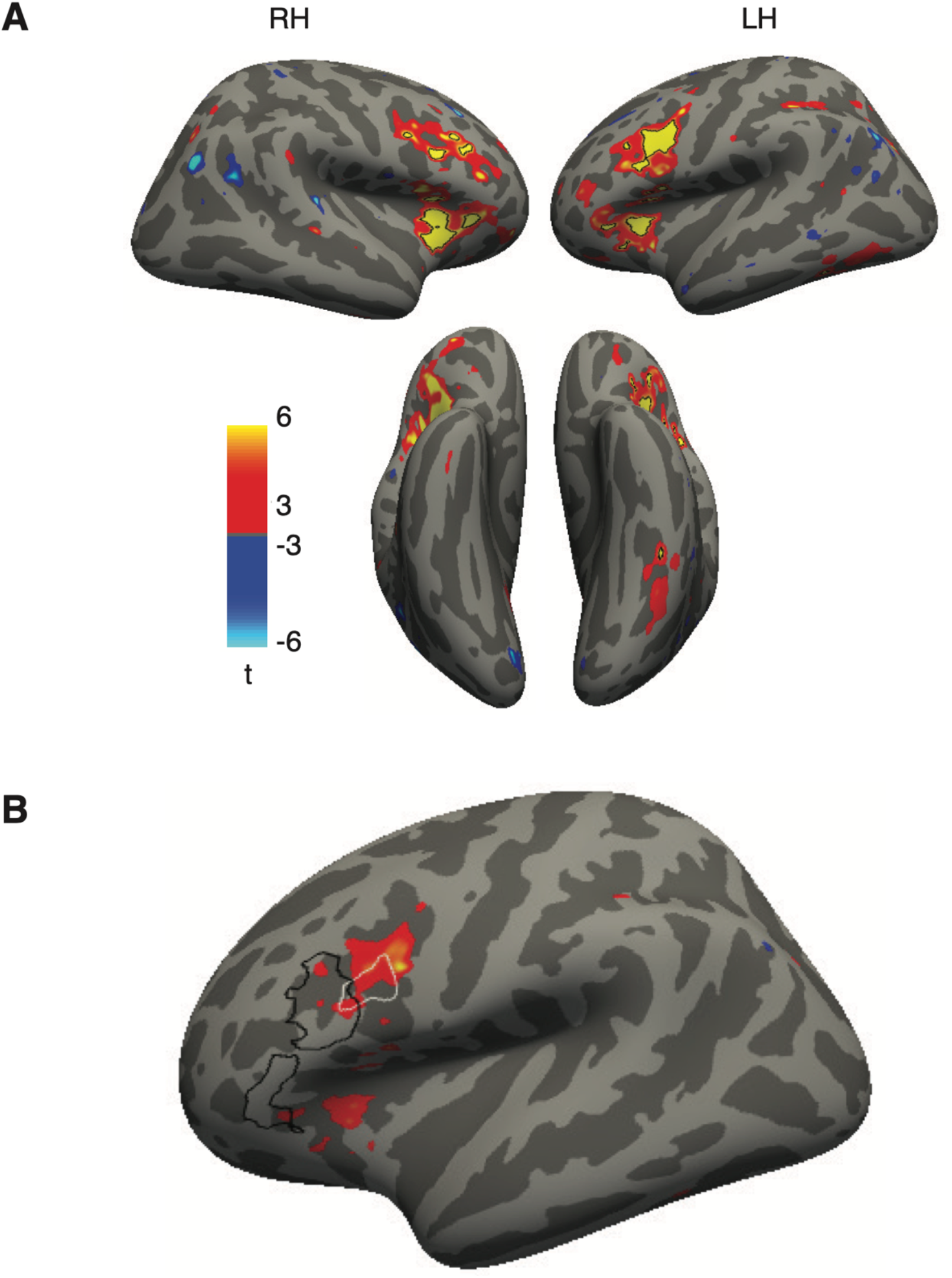
Whole-cortex analysis of the interaction between task and stimulus type. (**A)** Surface map (in the fsaverage template) of the interaction between stimulus type (text - false fonts) and task (lexical decision - fixation color). For each stimulus type, we computed differences in responses during the lexical decision task and the fixation color task. We then computed a difference of these difference maps for the two stimulus types. The resulting contrast map shows all nodes that had t-values > 3. Black outlines represent the patches with FDR-corrected p<0.05. **(B)** Lateral view of the left hemisphere, with the same interaction contrast as the colored blobs (corrected p<0.05). The outlines represent the ROIs analyzed in Figure 4: the two regions outlined in black together form “Language Broca’s area” and the more posterior area in whit is “Word Form Broca’s”.

Four regions demonstrated a significant interaction: (1) the left occipitotemporal sulcus, where the VWFA is located; (2) portions of the precental sulcus and inferior frontal sulcus (bilaterally), where, in the left hemisphere, our “Broca’s” ROIs are located; (3) the anterior insula / medial frontal operculum (bilaterally), and (4) a small portion of the left intraparietal sulcus (IPS). The first two of those regions overlap closely with our pre-defied ROIs (see **Figure 2** and **Figure 4**), although the right hemisphere homologue of ‘Broca’s area’ did not.

The latter two regions with a significant interaction were not pre-defined ROIs, but they have previously been implicated in reading. The anterior insula is often highlighted in fMRI studies of language function (Oh et al., 2014) although in some cases activity localized there may originate in the overlying frontal operculum (Fedorenko et al., 2015; Woolnough et al., 2019, 2023). The IPS region is also involved in cognitive processes relevant to reading (Rapp et al., 2016; Forseth et al., 2018; Woolnough et al., 2022) and is functionally correlated with the VWFA and Broca’s area (Vogel et al., 2012; White et al., 2023), as shown below.

### Modulations of VWFA activity are accompanied by task-dependent functional connectivity with the language network

To investigate how the areas that are modulated by reading task demands may be communicating with each other, we performed a functional connectivity analysis. Specifically, we correlated the *fluctuations* in trial-evoked responses, after subtracting out the mean response in each condition (White et al., 2023). For each surface vertex, we averaged the responses for each unique combination of task, stimulus type (real words, pseudowords and false fonts) and stimulus location. We then removed this mean response from the responses to all trials of the same unique condition, and performed the subsequent analysis with these demeaned *residual* responses. Specifically, we computed correlations coefficients in these residual responses across regions, separately for each condition.

**Figure 6A** shows a whole-brain map of correlations between each surface vertex and the left Language Broca’s area as the seed region. This represents functional connectivity during the lexical task (all stimulus types). Autocorrelation within the seed region during the lexical decision task was r=0.50. We observed positive correlations between the Language Broca’s area and much of the left superior temporal lobe, left intraparietal sulcus (IPS), and bilateral occipito-temporal sulcus, where the VWFAs are. There was also a hot spot of connectivity with a homologous ‘Broca’s area’ in the right frontal cortex. This map is consistent with prior results (White et al., 2023). **Figure 6C** shows analogous analyses with Word Form Broca’s area, VWFA-1, VWFA-2 and as the seed regions. The whole-brain maps of connectivity with the left VWFAs looked quite similar to the map in **Figure 6A** (with the seed in Broca’s area), but with somewhat less correlated activity in the superior temporal lobe. We also analyzed activity in anatomically-defined Brodmann’s areas 44 and 45, which are typically associated with “Broca’s area.” Activity there, in terms of BOLD responses and function connectivity patterns, resembled what we observed in “Word Form Broca’s.”

In addition to whole brain functional connectivity, we also computed mean correlations between the left VWFA and left Language Broca’s area, separately for each task and stimulus condition. For this analysis, we first averaged the single-trial beta responses across all nodes within each ROI, which reduces noise in the subsequent correlations. The results are in **Figure 6B**. Of primary interest here is how the functional connectivity between the VWFA and Broca’s area significantly *changed* depending on the task demands and stimulus content. We found a main effect of task (F(2,15)=21.63, p < 0.001) and stimulus type (F(2,15)=4.87, p = 0.03) on the correlation coefficients, and an interaction between them (F(2,396)=3.09, p = 0.04). We observed highest correlations between the two regions during the lexical decision task, for both types of stimuli, and lowest correlations during the fixation task. The correlation between these two regions in their responses to text were roughly twice as strong during the lexical task as during the fixation task. Statistics on these comparisons are reported in **Table 3**.

**Table 3:**
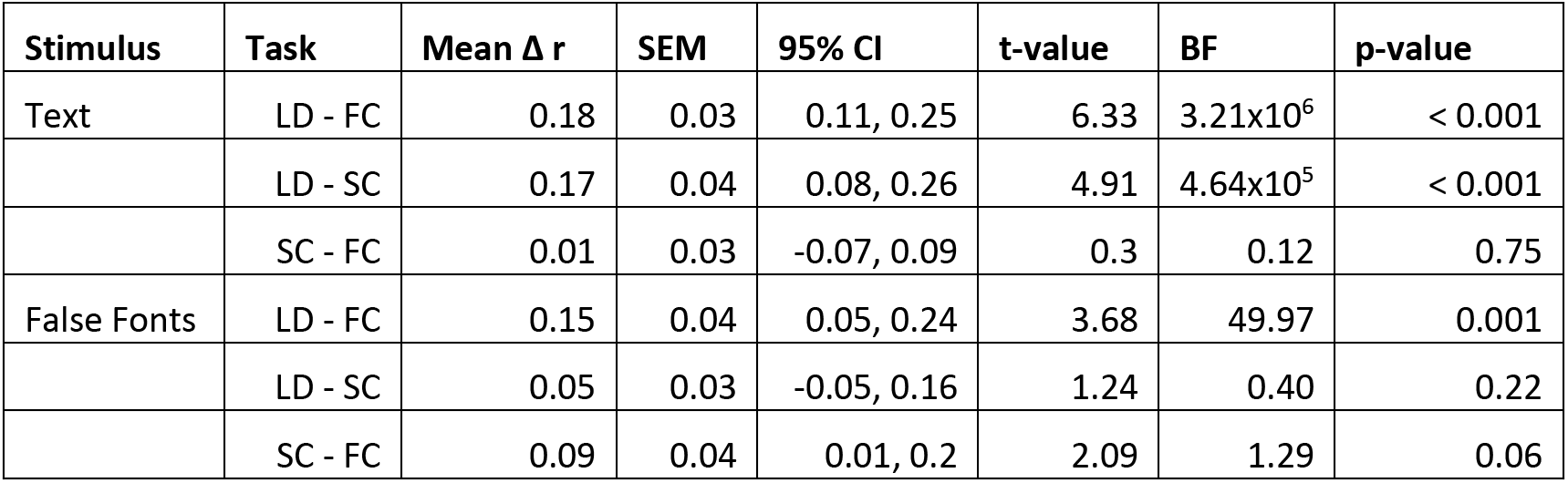
Functional Connectivity between Broca’s region and VWFA: Statistics on how functional connectivity between the language Broca’s Area and the left VWFA (collapsing over VWFA-1 and VWFA-2) differed across pairs of tasks. This is calculated separately for trials with text stimuli (both words and pseudowords) and false fonts. “Mean Δ r” is the mean difference in the correlation coefficient r across the two tasks listed in each row. P-values are FDR-corrected for multiple comparisons. Abbreviations: LD = lexical decision task; FC = fixation color task; SC = stimulus color task.

It is noteworthy that functional connectivity between the VWFA and Broca’s *increased* during the lexical task compared to the fixation task, even when false fonts were presented. This goes in the opposite direction as the mean overall response magnitudes, which were *decreased* for false fonts during the lexical task (**Figure 2A-B)**. This suggests that the suppression of response magnitudes to false fonts in the VWFAs could be caused by communication with Language Broca’s area.

### Task-driven correlations in BOLD responses and reaction times

Response times (RTs) are known to correlate with BOLD responses in several cerebral networks (Yarkoni et al., 2009; Domagalik et al., 2014; Mumford et al., 2024). The simplest explanation is that RTs indicate “time on task,” or the duration of time that the brain spends processing a stimulus. If the neural activity lasts for more time, the BOLD signal (which sluggishly integrates neural activity over time), yields a response of greater magnitude.

We conducted several analyses to determine whether a general effect of task difficulty on BOLD responses can explain the primary results reported above. First, we conducted a whole-cortex analysis of the correlation between BOLD response and RTs on single trials. The result is shown in the top row of **Figure 7A**: the mean correlation coefficients between normalized RTs and single-trial beta responses, for all stimulus types during the lexical decision task. These two measures were both de-meaned within each experimental condition (stimulus type and location) before entering into the correlation. The regions with significant correlation overlap partially with the regions highlighted as having functional connectivity with Broca’s area and the VWFA (**Figure 6A and 6C**), including left inferior frontal gyrus, insula/frontal operculum, pre-central sulcus, intraparietal sulcus, and occipito-temporal sulcus. In the right hemisphere (not shown) during the lexical decision task we observed hotspots of correlations with RT in the insula/frontal operculum and anterior cingulate cortex.

**Figure 6:**
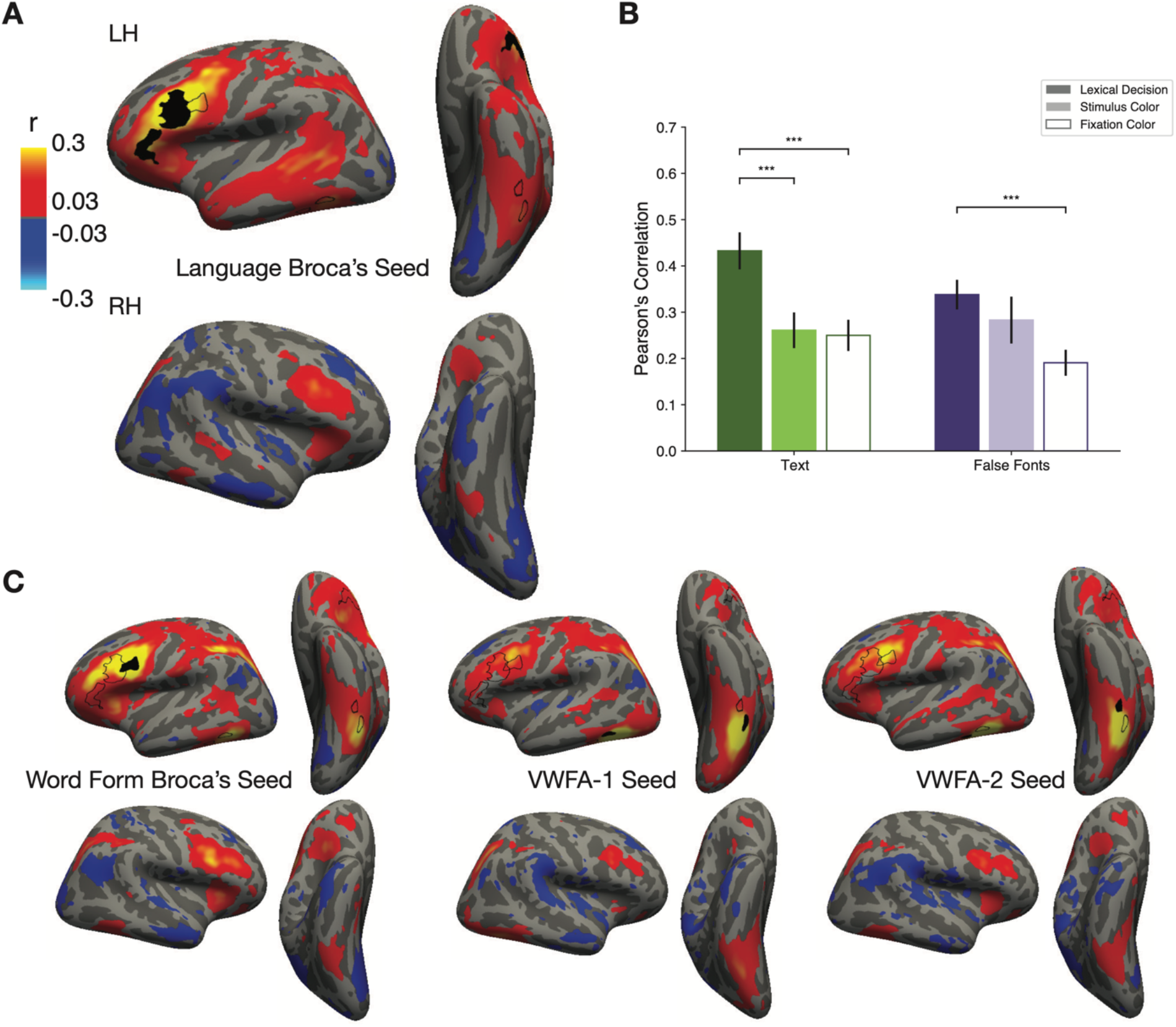
Functional connectivity shows task-dependent communication between Broca’s area and the VWFA. **(A)** Whole-brain connectivity with the seed region in left language Broca’s area (black outline) during the lexical decision task for all stimulus types. The color map represents Pearson’s correlation coefficients (minimum r>0.11, p<0.05, FDR corrected). The two black outlines on the left ventral surface represent the peaks of the contrast between words and all other categories in the category localizer experiment, in the average brain. Thus, they roughly correspond to the most likely locations of VWFA-1 and VWFA-2. (**B**) Average correlation coefficients between left language Broca’s area and the left VWFA (collapsing VWFA-1 and VWFA-2). We first averaged responses for each condition within each of the two ROIs, de-meaned them and then computed pairwise correlation coefficients. The correlations were highest during the lexical decision task for both types of stimuli. Error bars are +/- 1 SEM. Horizontal lines and stars indicate statistical significance. (*=p<0.05, **=p<0.01, ***=p<0.001). **(C)** Correlation maps like in panel A, but for three different seed regions: Word Form Broca’s Area, left VWFA-1, and left VWFA-2.

The bottom row of **Figure 7A** is the same analysis but for trials during the fixation task, when the same stimuli were presented and participants also had to make a response on each trial. Compared to what was observed during the lexical task (top row), the correlations between BOLD and RT were markedly weaker in the vicinities of Broca’s area and the VWFA (left inferior frontal sulcus and occipito-temporal sulcus). Thus, to the extent that there is a relation between the difficulty of each trial and the activity in these regions, it is not domain-general but rather specific to the task that requires reading the words.

**Figure 7:**
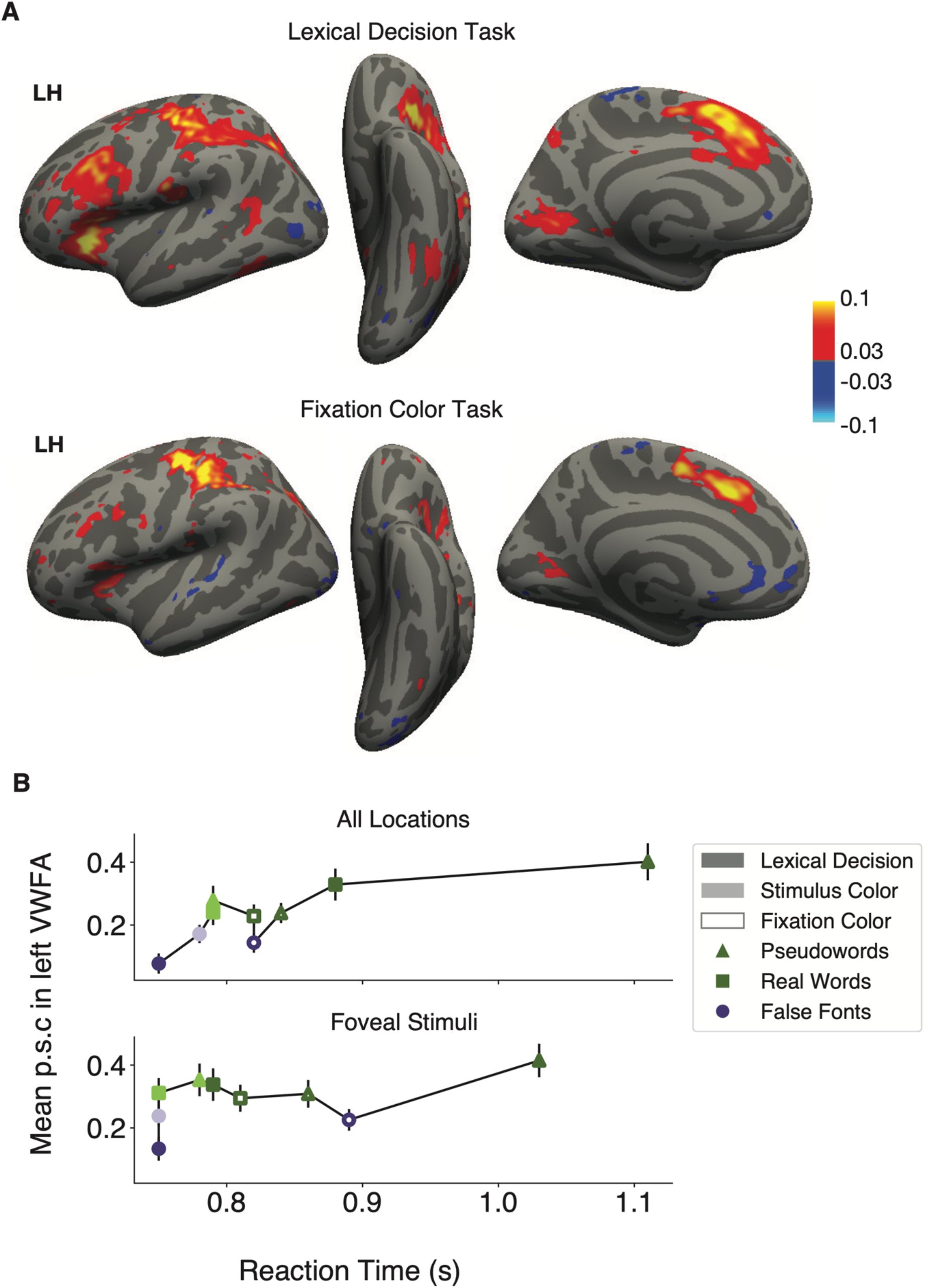
Relationship between reaction times and BOLD responses. **(A)** Whole-cortex correlations between response times and magnitude of BOLD response. The color map represents Pearson’s correlation coefficients (minimum r>0.03, q<0.05). The data in all panels corresponds to trials with correct behavioral responses only. The top panel shows correlations for the lexical decision task, and the lower panel for the fixation color task. **(B)** Mean left VWFA response magnitudes as a function of mean response times in each stimulus and task condition (correct trials only). Error bars represent ±1 SEM. The top panel averages over all stimulus locations, and the bottom is only for trials with stimuli at the central (foveal) location.

The second analysis concerns the relation between RTs and activity in the left VWFA specifically, across all conditions. **Figure 7B** plots the mean BOLD response in the left VWFA as a function of the mean RTs in each condition, on correct trials only. The top graph is for all stimulus locations, and the bottom graph is for trials when the stimuli were presented at the fovea (0°). The three stimulus types are represented by different shapes, and the three tasks are represented by different shadings. In the top graph, there is clearly a positive correlation between RT and BOLD. This may provide an explanation for the VWFA’s responses during the lexical decision task: false fonts have the smallest RTs and smallest BOLD responses, whereas pseudowords have the largest RTs and largest BOLD responses. This could be because when a false font string is presented, the participant, and their VWFA, quickly recognizes that it is not a word and stops processing it. In contrast, pseudowords look plausibly like real words, so the brain must effortfully search through the mental lexicon before deciding to reject the stimulus. Thus, correct RTs are longer and the BOLD response is higher for pseudowords, compared even to real words, which can be identified relatively quickly (Woolnough et al., 2021; Gagl et al., 2022). However, Taylor et al. (2013) demonstrated that even when regressing out the effect of RT, pseudowords still evoked a larger response in the VVFA than real words.

We also used a model comparison approach to demonstrate that RTs, by themselves, do not explain all the variance in the BOLD responses. We first constructed a model that included only response times as a fixed effect predictor, with random effects of response time, stimulus type and task across subjects. We then constructed a second model with the same random effects, but added in stimulus type and task and their interactions as additional fixed effects. We found that the second model fit the VWFA better than the first (𝛘^2^(10) =112.64, p<0.001), even when penalizing the second model for its greater number of parameters (BIC 9268.3 for the second model compared to 9289.2 for the first; lower BIC values indicate better model fit). The weaker fit of the RT-only model is made clear by how the VWFA responds during the fixation color task (open symbols in **Figure 7B**). During this task, there is no clear relationship between RT and BOLD magnitude across stimulus types. Moreover, RTs were generally longer in the fixation task than the stimulus color task, but BOLD response magnitudes were similar between them.

Also, if we consider only trials when the stimulus was presented at fixation (excluding stimuli 3° to the side), the influence of RT is reduced but the task and stimulus effects are still strong. That is shown in the lower plot in **Figure 7B**. Again, the model that included task and stimulus type as predictors in addition to reaction time fit the data better than one that only included reaction times (𝛘^2^(10) =40.82, p<0.001; BICs 3117.2 vs. 3138.0, respectively). Thus, we conclude that much of the shared variance between RTs and BOLD that produced the pattern the upper plot was explained by the increased difficulty when the stimuli are in the parafovea. Without that difficulty, BOLD is less related to RT but still strongly influenced by the interaction of task and stimulus.

## Discussion

We investigated the combination of stimulus features and task demands that maximally engage the VWFA. Consistent with the hypothesis that this region is a module tuned for a special class of stimulus, we found that it responds more strongly to words than strings of unfamiliar characters even during a demanding visual task that withdraws attention from the stimuli. Nonetheless, the results show that the VWFA’s activity can neither be solely accounted for by stimulus selectivity, nor by top-down modulation. The strongest activation requires specific stimuli (strings of familiar letters) presented during specific tasks (explicitly trying to read words). Engaging in the lexical task increased the VWFA’s BOLD response to words and pronounceable pseudowords (with an even greater increase for pseudowords), but *suppressed* the response to visually-matched unfamiliar characters (false fonts). Thus, top-down language processing does not simply *add* to the VWFA’s BOLD activity; rather, the modulation differs greatly across types of character strings. Moreover, compared to when the stimuli were ignored, attending to either type of character string to judge its color had little impact on the VWFA’s response.

This interaction between bottom-up and top-down factors was unique to the VWFA among visually-driven areas. Given the patterns of functional connectivity, which also differed greatly across tasks, we suggest that the modulations targeted to the VWFA are due to interactive feedback from regions involved in processing language.

### Possible explanations for what drives activity within the VWFA

By fully crossing task demands and stimulus types, our results rule out four hypotheses about what determines the magnitude of VWFA responses:

*(1) Word-likeness of visual stimuli:* One hypothesis is that the VWFA’s activity is explained by bottom-up tuning for visual stimuli that are the most like known words. That predicts, in our study, that responses should be weakest for false fonts, intermediate for pseudowords, and highest for real words, in all three tasks. Our data only partly match that prediction. The VWFA did prefer real letter strings over false fonts across all tasks. But that selectivity was greatly magnified during the lexical task (compared to both other tasks). More importantly, the stronger response to pseudowords than real words occurred only occurred during the lexical decision task (**Figure 2C**). These results suggest that the VWFA’s sensitivity to *lexical* attributes of letter strings is contingent on top-down cognitive influences during linguistic tasks.
*(2) Top-down boost for attended stimuli:* Throughout visual cortex, task-relevant stimuli evoke larger responses than task-irrelevant or ignored stimuli. These attentional effects could be described as additive or multiplicative modulations of the stimulus-driven response, with a magnitude that depends on the degree of similarity between the tuning properties of neurons and what is attended (Reynolds and Heeger, 2009). A related hypothesis for the VWFA is that attentional mechanisms boost responses to task-relevant stimuli, but only when the stimuli are text. That hypothesis would predict a higher response to words when they are task-relevant than when they are ignored, and no modulation of the response to false fonts.
Our data violate this prediction in two ways. First, the VWFA responded similarly to attended words during the stimulus color task and to ignored words during the fixation color task (light green bar in **Figure 2B**). Second, the response to false fonts was *lower* when they were task-relevant during the lexical decision task than when they were ignored (dark purple bar in **Figure 2B**). Together, these results show that the top-down modulations in the VWFA cannot easily be explained by visual attention, consistent with White et al. (2023).
*(3) Automaticity of word recognition:* This hypothesis implies that as long as a word is legible and some attentional resources are available, its meaning is fully processed (Carr, 1992; Brown et al., 2002). In its simplest form, such a theory would predict equivalent responses to written words in the lexical decision and stimulus color tasks. In contrast, we found that engaging in the stimulus color task had a negligible effect compared to the fixation task. Therefore, strong VWFA activation requires voluntary effort to read the words.
*(4) Task difficulty:* This hypothesis is that the VWFA, along with other regions, responds more strongly on trials when the participant makes a more difficult judgment that requires more processing time. Several studies of word reading have found correlations between the behavioral RTs and BOLD magnitudes in the VWFA as well as a wider network of regions (Binder et al., 2005; Graves et al., 2010; Taylor et al., 2013).

In our data, two patterns suggest that this hypothesis cannot fully explain the modulations of VWFA activity. First, in the VWFA and Broca’s area, the correlation between RT and BOLD is not a *general* effect of the effort required on each trial. Rather, it is much stronger during the lexical decision task than the fixation task (see the maps in **Figure 7A**). Second, VWFA responses are best predicted by a model that includes task and stimulus conditions in addition to mean RTs (see **Figure 7B**). Nonetheless, during the lexical decision task there is a relation between BOLD and RT, in terms of the differences between pseudowords, real words, and false fonts. We believe that this is due to a top-down signal that prolongs the VWFA’s processing of pseudowords and shuts down processing of false fonts.

### Our proposal for the role of the VWFA and its functional connectivity

Our results support the theory that the VWFA is primarily a visual region devoted to identifying familiar letter strings (Vinckier et al., 2007; Dehaene et al., 2015). Its bottom-up responses are selective for familiar letter strings, even when attention is withdrawn. However, the VWFA is not automatically engaged in word recognition when familiar letter strings are presented and attended. Its selectivity for text is greatly enhanced when the task requires reading, in part due to active suppression of non-letter stimuli. Also, its sensitivity to lexical features of stimuli depends critically on the participant engaging in an explicitly linguistic task.

These aspects of its function are dependent on communication with other parts of the language network, as our functional connectivity analyses revealed (**Figure 6**; see also (Vogel et al., 2012; Stevens et al., 2017; Chen et al., 2019; López-Barroso et al., 2020). We also discovered higher correlations between the VWFA and Broca’s area during the lexical decision task than the other two tasks. This task-dependent modulation of connectivity was unique to the VWFA within ventral temporal cortex (White et al., 2023). Other researchers have also reported that the connections between VWFA and frontal regions change as a function of familiarity with a language (Boukrina et al., 2014).

Importantly, correlated activity with Broca’s area was strong both when the VWFA’s response magnitudes were enhanced (for text) and when they were suppressed (for false fonts during the lexical decision task). The mean BOLD responses in the seed region used for this analysis, which we called Language Broca’s area, offer clues to the nature of its role. Its BOLD signal deviated from baseline only during the lexical decision task, with a positive response to letter strings and a negative response to false fonts (**Figure 4A**). This suggests that the suppression of the VWFA response to false fonts during the lexical decision task is due to top-down feedback from Broca’s area.

### Limitations and future directions

Methods that offer greater temporal resolution, such as magnetoencephalography or intracranial recordings, will be necessary to reveal *when* the task effects arise during the processing of a word. Past research gives some clues: intracranial recordings from ventral temporal cortex showed that responses to attended words differed from ignored words relatively late, ∼400-800 ms after word onset (Nobre et al., 1998). However, the time course of attention effects may also depend on the task, as engaging in an orthographic task has also been shown to modulate an earlier evoked potential (the N200; Ruz and Nobre, 2008).

By using fMRI, we were able to demonstrate the spatial specificity of cognitive modulations in the VWFA, which did not occur in nearby category-selective regions. Nonetheless, there may be analogous cognitive effects in those other regions during tasks optimized for them, such as face recognition in the FFA.

Lastly, there is more to learn about differences between word-selective sub-regions. We found similar activity patterns in VWFA-1 and VWFA-2. While some previous studies have reported the same (White et al., 2023), others have emphasized the importance of functional subdivisions of the VWFA (Lerma-Usabiaga et al., 2018). Similar questions remain about sub-regions in the vicinity of “Broca’s area.” Other researchers have speculated about functional segregation within this general region along the dorsal-ventral axis, as well as the anterior-posterior axis (Hagoort, 2014).

### Conclusions

We conclude that activity in the VWFA is determined by an interaction between inherent stimulus selectivity and top-down input from frontal language regions. The VWFA is maximally activated when letter strings are presented *and* the observer is voluntarily looking for known words. It is not sufficient for words to be presented or even attended visually. Moreover, engagement in a lexical task does not always enhance the VWFA’s response; in fact, it suppresses the response to unfamiliar characters. These findings open the door to future work on more naturalistic reading tasks, as well as comparisons to other specialized parts of the visual system.

## Acknowledgments

We are grateful to Nicole Oppenheimer and Kimya Firoozan for help with data collection, Oscar Woolnough for advice on the insula, and to the National Eye Institute for funding (grant R00 EY-029366).

## Extended Data

**Extended Data Figure 1-1:**
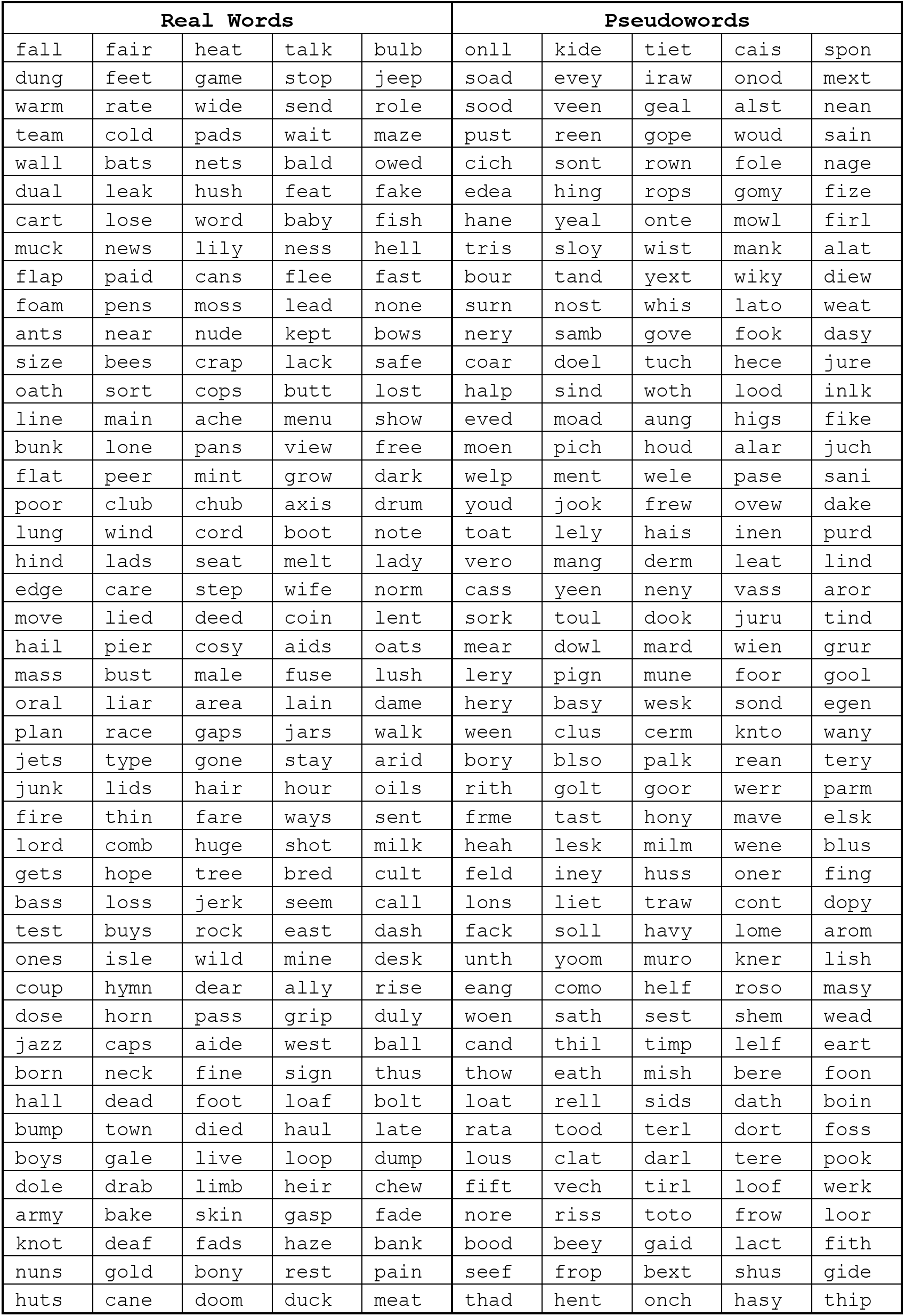
All words and pseudowords used in the experiment.

**Extended Data Figure 1-2:**
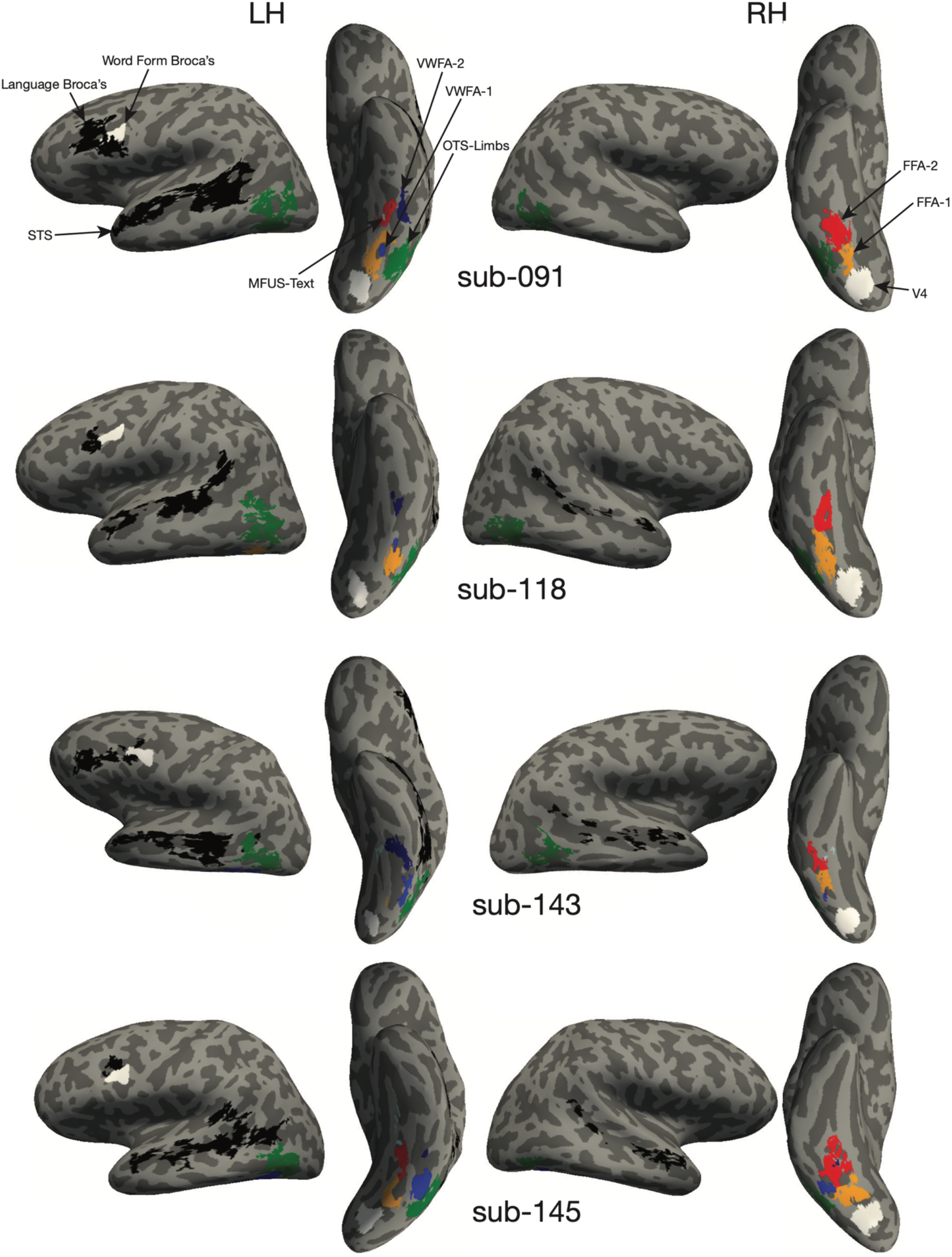
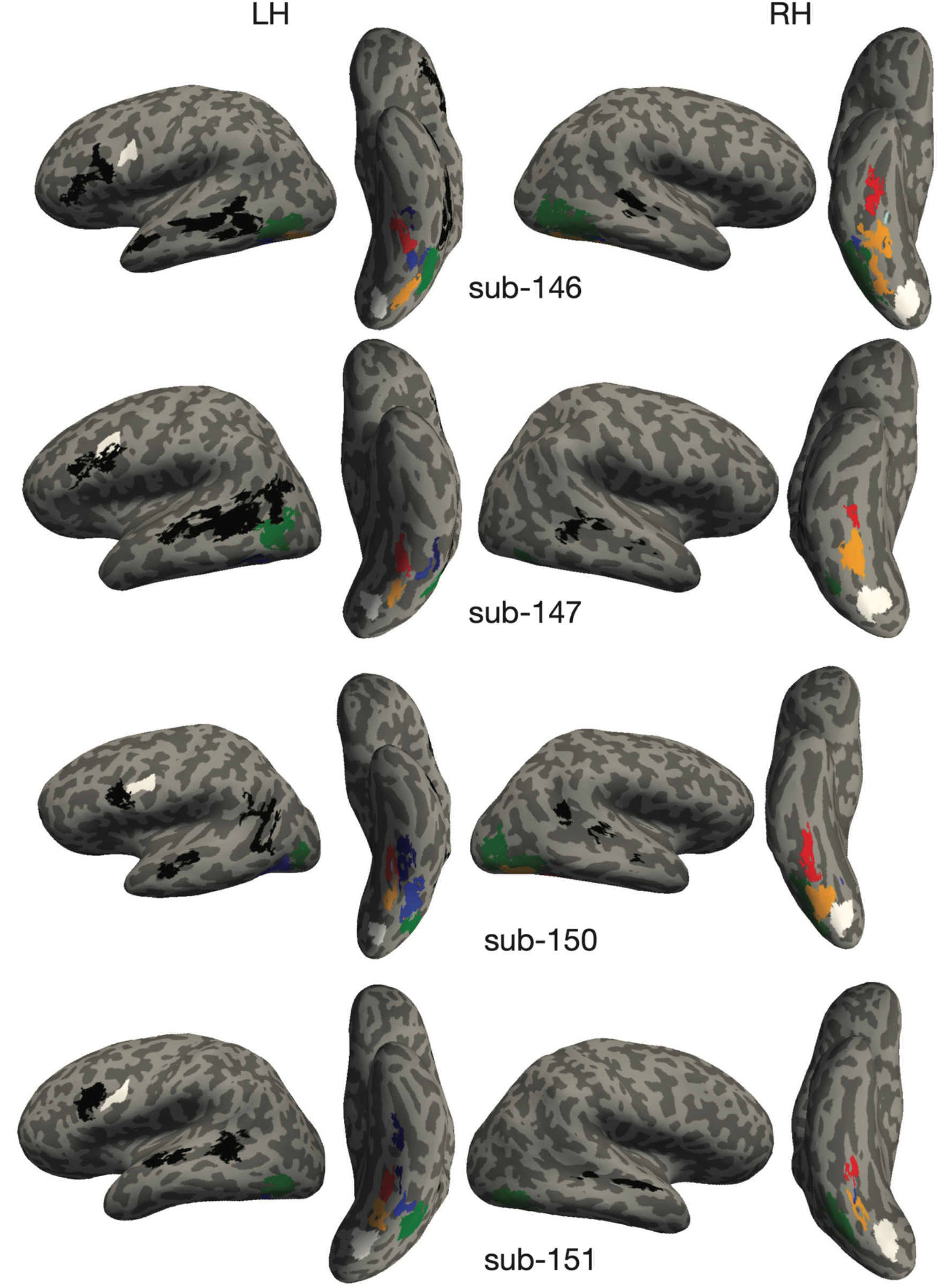
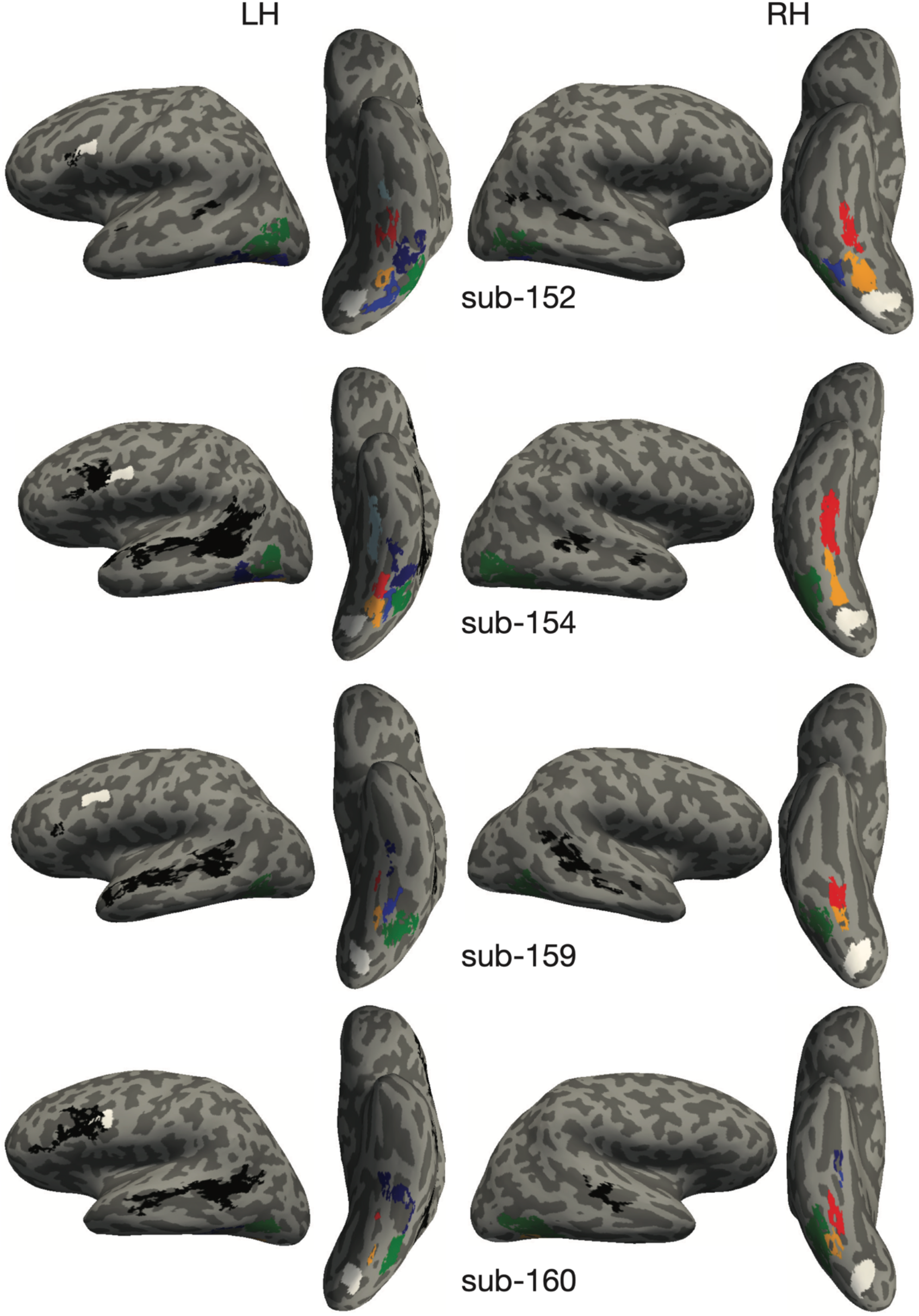
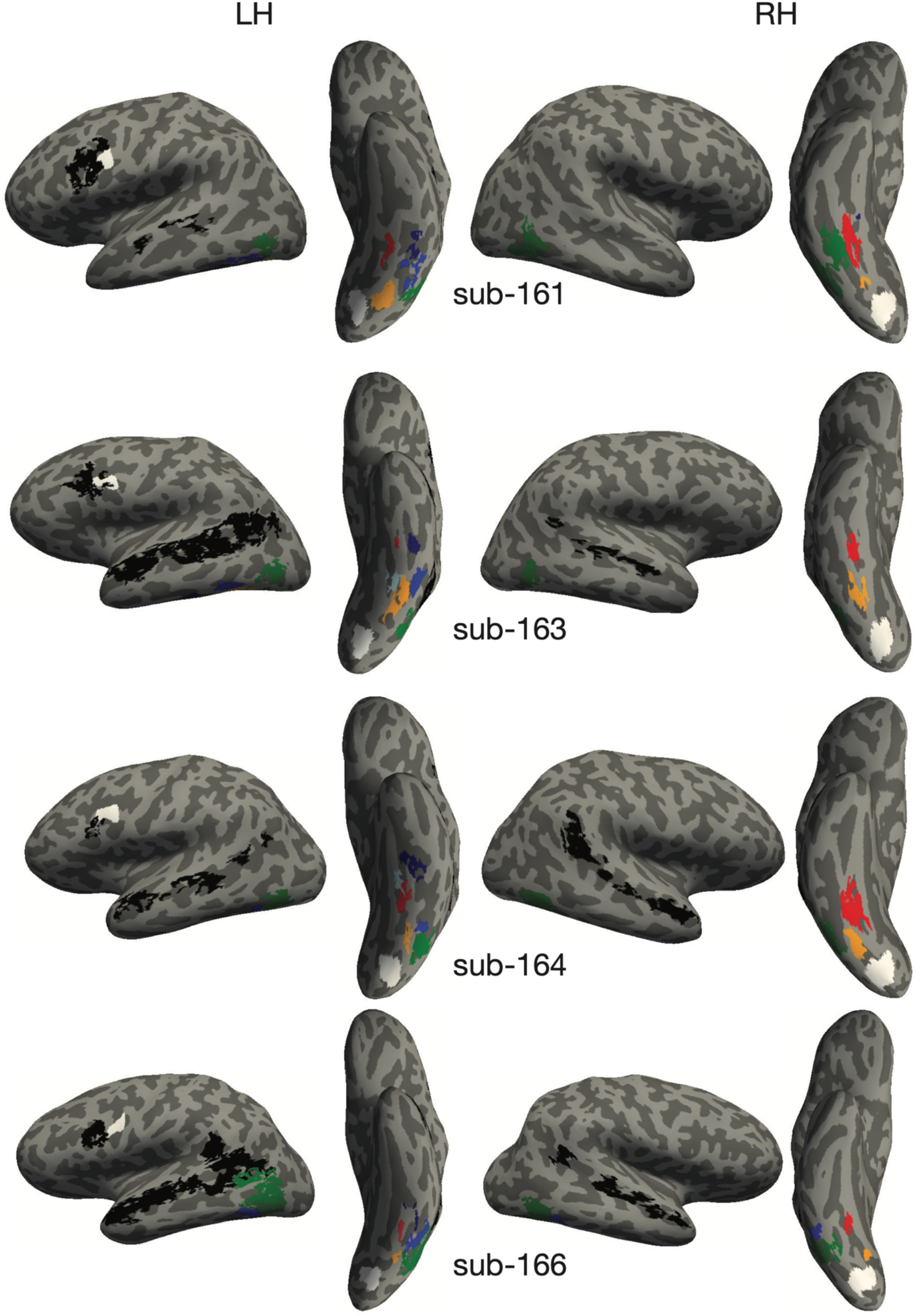
Regions of interest in individual subject anatomical surface space.

**Extended Data Table 2-1:**
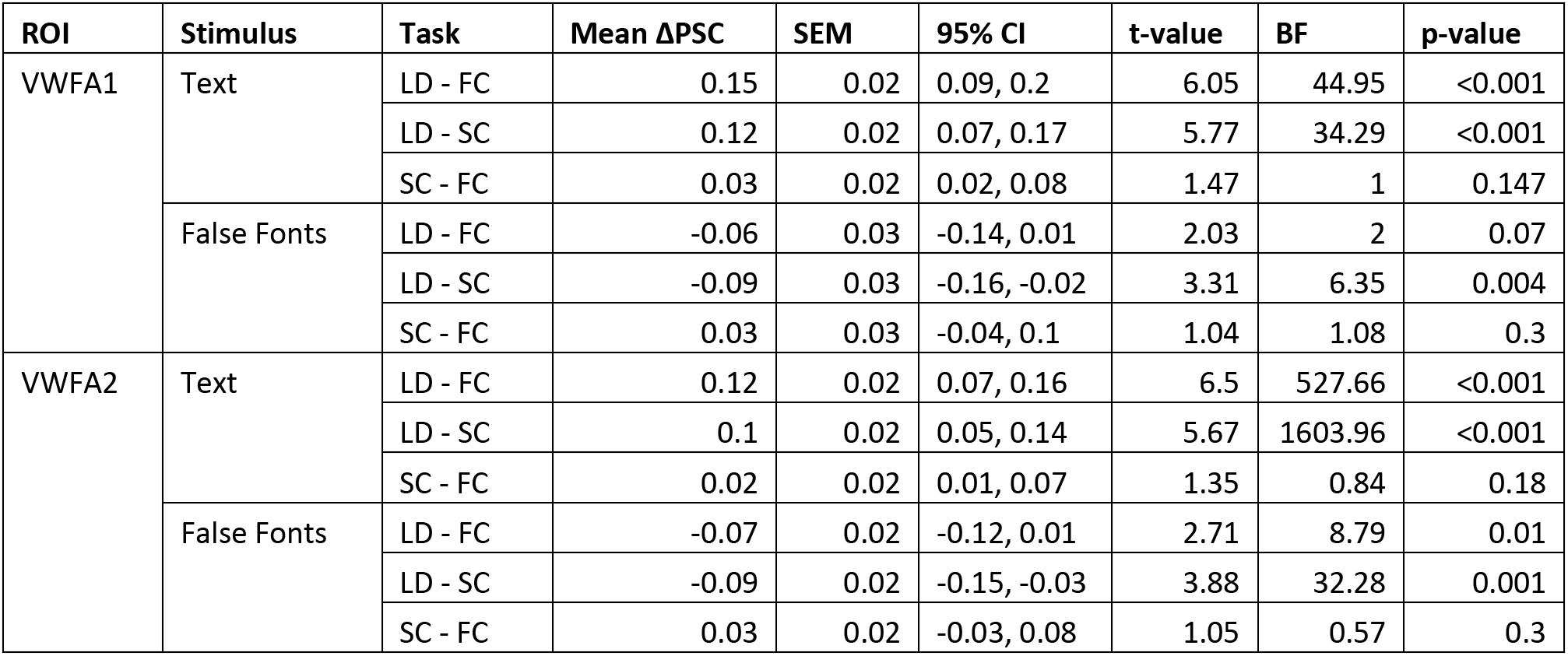
Statistics for stimulus and task differences for VWFA-1 and -2, reported separately.

**Extended Data Figure 2-1:**
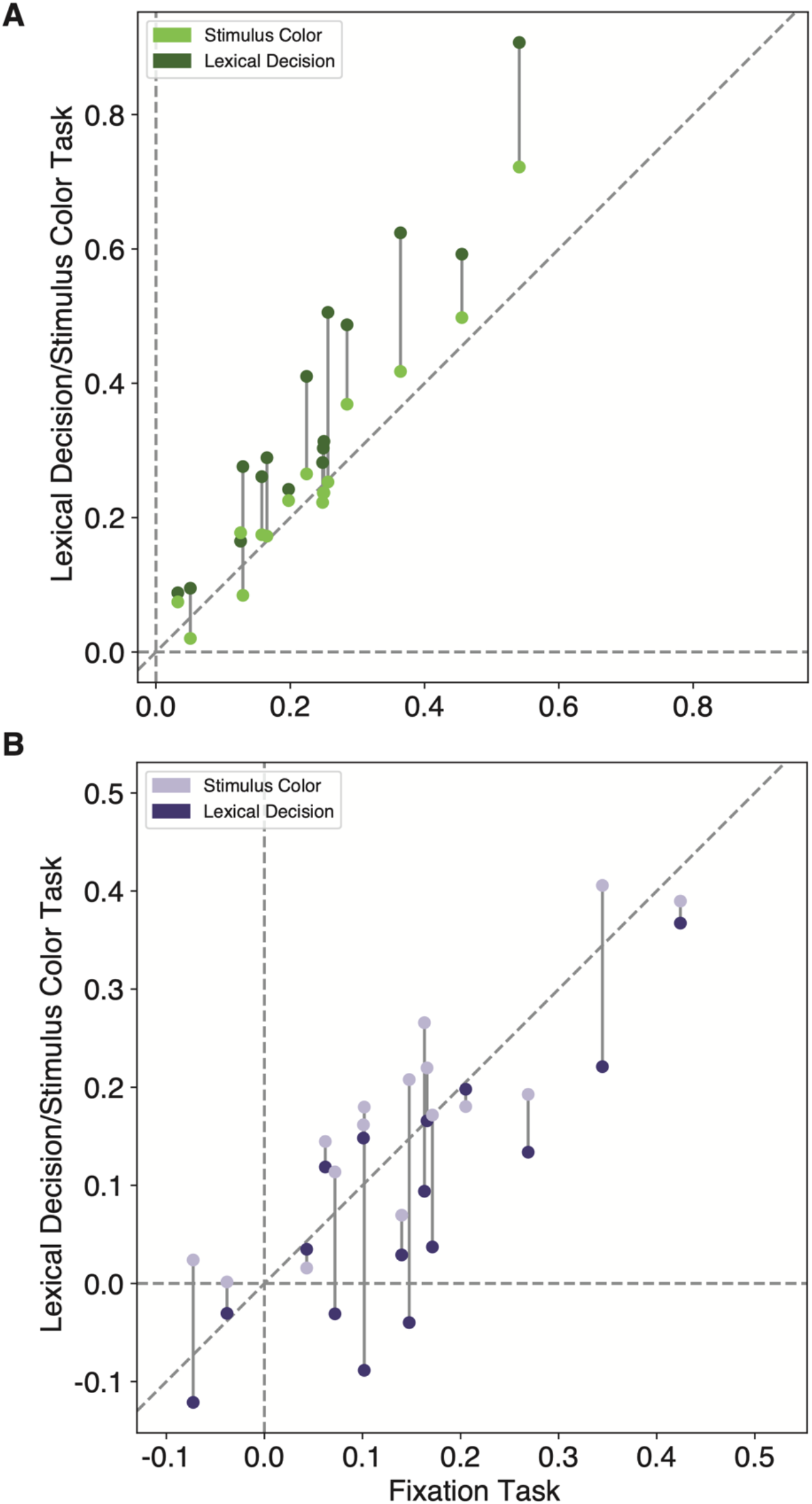
Individual participants’ mean beta responses in the left VWFA (VWFA-1 and VWFA-2 together) for **A)** text (real words and pseudowords) and **B)** false fonts. In each panel, each participant contributes two points that are connected by a vertical line. The horizontal position of each point represents that subject’s VWFA response during the fixation task. The vertical position of the darker points represents the response during the lexical decision task, and the vertical position of the lighter points represents the stimulus color task.

**Extended Data Figure 3-1:**
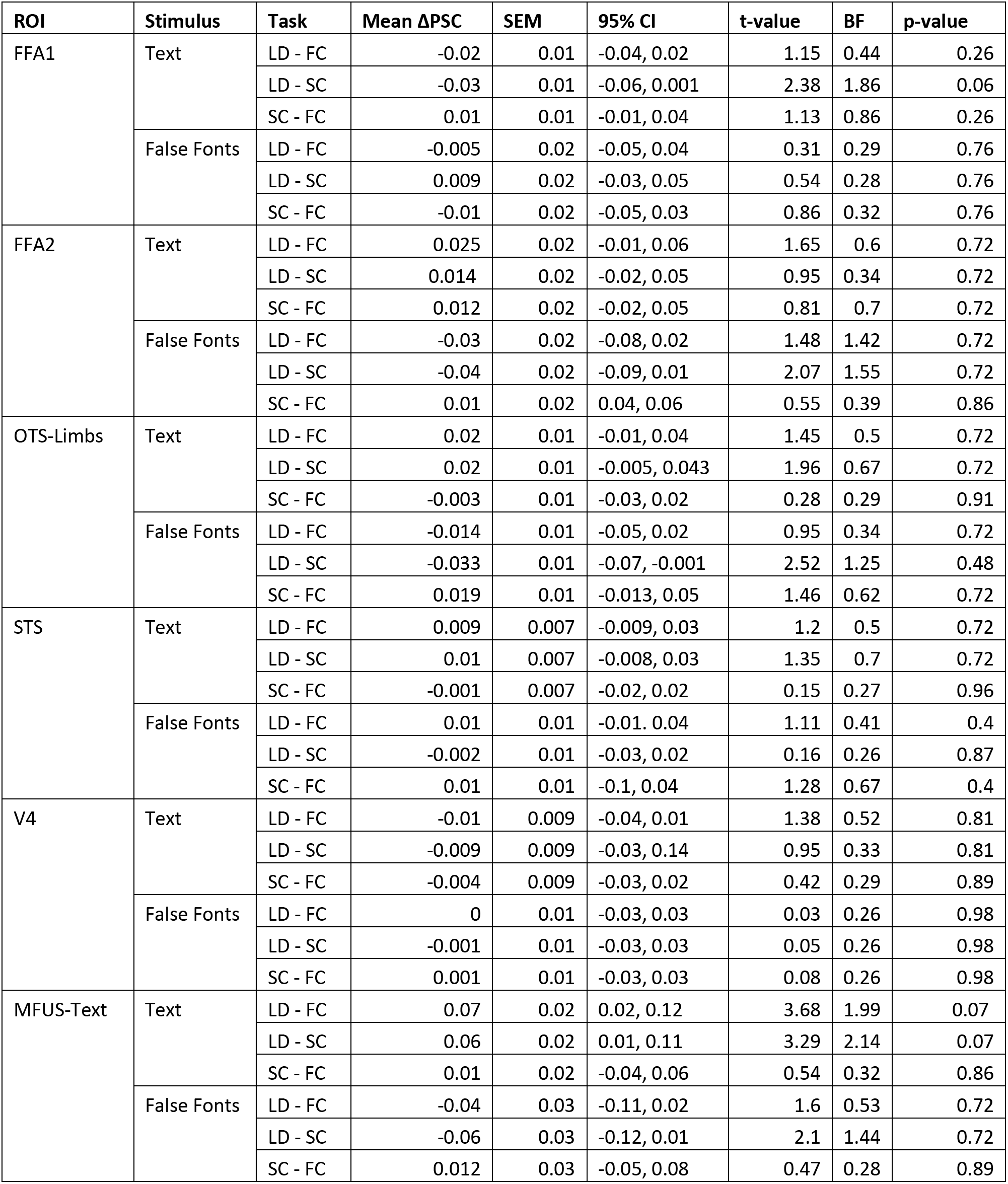
Statistics for task effects on responses to each stimulus type for all control ROIs in the left hemisphere. P-values are corrected for multiple comparisons using FDR correction. All p-values were corrected for post-hoc tests within ROIs, and then concatenated and corrected for multiple comparisons across ROIs. Adjusted P-values are reported in the final column of the table.

**Extended Data Figure 3-2:**
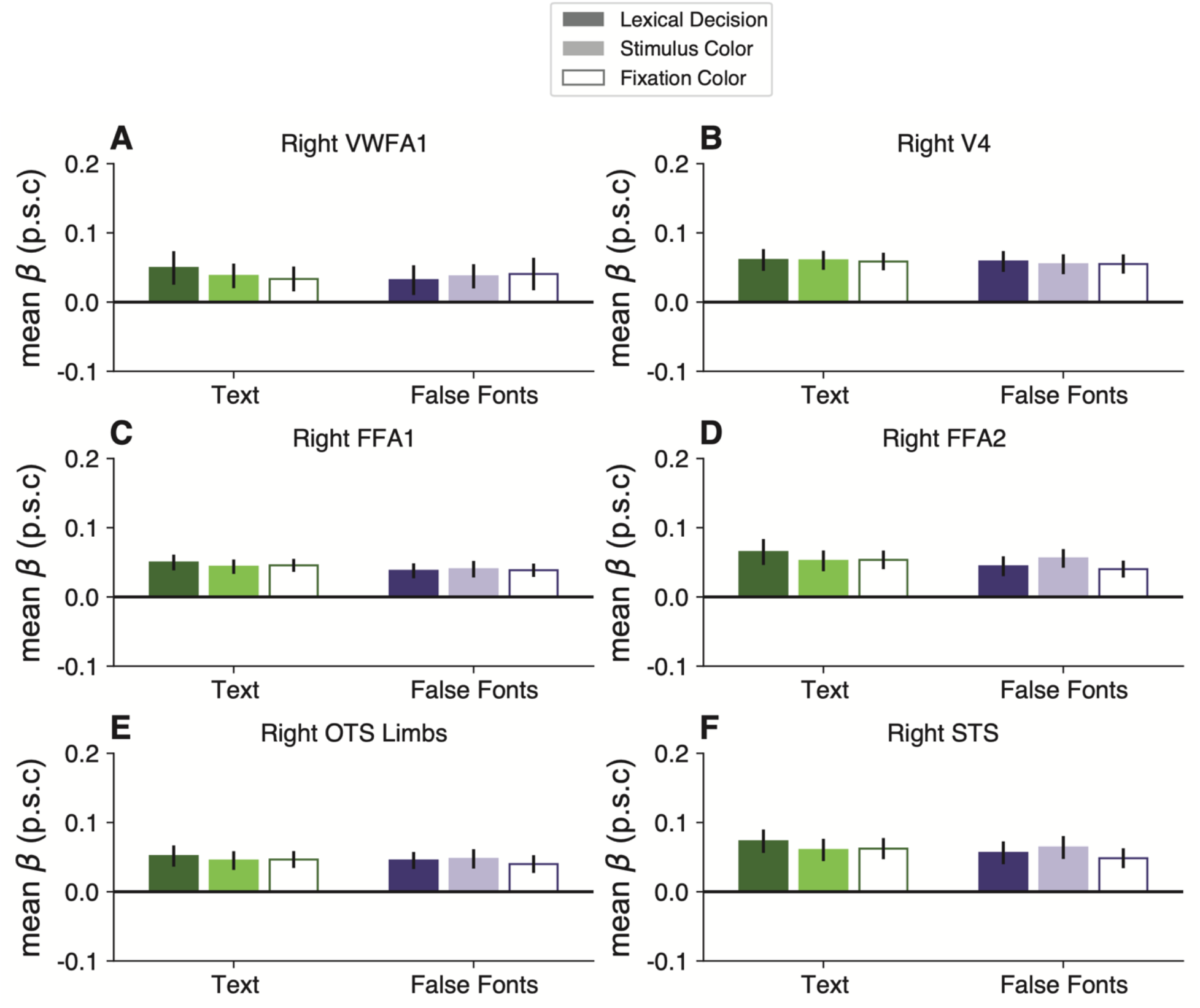
Mean BOLD percent signal change as a function of stimulus and task types in control ROIs in the right hemisphere. Error bars represent standard error of the mean.

